# Recreational climbing alters cliff soil chemistry and plant-associated fungal communities

**DOI:** 10.64898/2026.05.15.725402

**Authors:** Ana García-Muñoz, Franz-Sebastian Krah, Gemma Palomar, Álvaro López-García, Mateusz Buczek, Juan Lorite, Martí March-Salas

**Affiliations:** Area of Biodiversity and Conservation, Department of Biology, Universidad Rey Juan Carlos-ESCET, Calle Tulipán s/n, 28933 Móstoles, Madrid, Spain; Instituto de Investigación en Cambio Global (IICG), Universidad Rey Juan Carlos, Calle Tulipán s/n, 28933 Móstoles, Madrid, Spain; Department of Plant Biology and Ecology, Universidad de Sevilla, Sevilla, Spain. Calle Reina Mercedes 6, 41012 Sevilla, Spain; Fungal Ecology and BayCEER, University of Bayreuth, Bayreuth, Germany. Universitätsstr. 30, 95447 Bayreuth, Germany; Global Change Research Institute of the Czech Academy of Sciences, Brno, Czech Republic. Bělidla 986/4a, 603 00 Brno-střed, Czech Republic; Department of Genetics, Physiology and Microbiology, Faculty of Biological Sciences, Universidad Complutense de Madrid, 28040 Madrid, Spain; Institute of Environmental Sciences, Faculty of Biology, Jagiellonian University, 30-387 Kraków, Poland; Soil Microbiology and Symbiotic Systems Department, Estación Experimental del Zaidín, CSIC, Profesor Albareda 1, 18008 Granada, Spain; Department of Botany. University of Granada (UGR). Faculty of Sciences. Avenida de Fuente Nueva, s/n, 18071 Granada, Spain; Plant Evolutionary Ecology, Institute of Ecology, Evolution and Diversity, Faculty of Biological Sciences, Goethe University Frankfurt, Max-von-Laue-Str. 13, 60438 Frankfurt am Main, Germany

**Keywords:** Arbuscular mycorrhizal fungi (AMF), chemical impact, edaphic specialization, extreme environments, human-related disturbances, plant-microbial interactions, recreational rock climbing, rhizosphere-associated symbionts

## Abstract

- Cliffs are environmentally extreme yet biodiversity-rich ecosystems that harbour specialist plants, many endemic and threatened. Plant persistence in these nutrient-poor substrates may depend on tightly linked soil- and root-associated microbial communities, which remain poorly understood. These interactions may become increasingly important with the global expansion of recreational climbing. While physical climbing impacts on vegetation are documented, potential chemical effects, from the use of climbing chalk (magnesium carbonate), on soil properties and plant-associated microbiota remain unknown.
- We sampled soils and roots beneath cliff-specialist and generalist plants, and unvegetated soils, across climbed and unclimbed routes in northern, central, and southern Spain. Soil physicochemical properties were quantified, fungal communities were characterized using ITS-metabarcoding, and structural equation modelling was used to disentangle direct and indirect effects.
- Climbing increased soil pH and altered soil chemical properties, driving shifts in fungal diversity and functional composition in soil and roots. The relative read abundance of root-associated symbiotrophic fungi declined, whereas arbuscular mycorrhizal fungi and pathogens increased in climbed cliffs. Overall effects were consistent, with cliff-specialist plants mediating nutrient and fungal shifts.
- ur findings show that climbing can reshape cliff soil chemistry and fungal communities, with potential cascading consequences for plant functional performance, nutrient dynamics, and ecosystem resilience.

## Introduction

Extreme ecosystems are widely recognized as key reservoirs of unique biodiversity, acting as refugia from various biotic and abiotic pressures and fostering the specialization of highly adapted plant lineages (Harrison & Noss, 2017). Harsh abiotic conditions, including edaphic restrictions, extreme temperatures, intense radiation, strong winds, limited water availability, and physical inaccessibility, apply strong environmental filtering, but at the same time, allow stress-tolerant specialized plants to persist and diversify (Harrison & Noss, 2017; Trew & Maclean, 2021). Under such environmental constraints, the persistence of specialized plants is often tightly linked to microbial associations, which can enhance nutrient acquisition, drought resistance, edaphic stress tolerance, and pathogen defence, ultimately helping ecosystem stability (Kohler et al., 2015; Liu et al., 2022). Though, microbial research remains underrepresented in most extreme systems, including cliffs, which limits our understanding of their ecological role.

Cliffs represent clear examples of ecosystems characterized by extreme environmental, physical and edaphic limitations for plant development and persistence (Larson et al., 2000; Fitzsimons & Michael, 2017). Soil development is minimal or absent and is primarily limited to accumulated substrate, water retention is low, and nutrient inputs can be limited and spatially heterogeneous (Larson et al., 2000; Langevin et al., 2024). Despite these constraints, cliffs frequently emerge as biodiversity hotspots, harbouring between 35-66% of the endemic flora in many countries, with numerous lineages specialized to these habitats for millennia (Larson et al., 2000; March-Salas et al. 2023a). Indeed, cliffs rank among the global systems with the highest proportion of threatened plant taxa (*c.* 46%; Nyberg et al., 2025). However, despite their ecological and conservation importance, the study of microbial communities associated with cliff plants remains anecdotal (but *see* Krah & March-Salas, 2021; Dumas et al. 2025; reference therein Zhao et al. 2025).

Nowadays, cliff ecosystems face two major and sometimes interacting threats. First, climate change is intensifying the frequency and duration of droughts, exacerbating water and nutrient limitations in systems already constrained by extreme abiotic conditions. Second, recreational rock climbing has experienced rapid expansion worldwide, with annual growth rates reaching up to 20% in some areas, particularly in Mediterranean regions, characterized by suitable topography and year-round favourable climates (deCastro-Arrazola et al., 2021). While the physical disturbance of cliff vegetation by climbing has been documented, including trampling, mechanical damage, and changes in plant cover (*e.g.*, Lorite et al., 2017; Schweizer et al., 2021; March-Salas et al., 2018, 2023b; Morales-Armijo et al., 2023), impacts of climbing activity on soil- and plant-associated microbial communities remain largely unexplored.

A central yet understudied factor of climbing disturbance is the widespread use of climbing chalk, primarily composed of magnesium carbonate (MgCO₃). As an inorganic compound, climbing chalk can accumulate in rock fissures and microsites where soil particles and organic matter are retained (Hepenstrick et al., 2020; Schweizer et al., 2021; Harrison et al., 2024). Its deposition may generate chemical alterations, potentially affecting nutrient composition and their physicochemical properties. An experimental study using Petri-dish assays has shown that climbing chalk can reduce germination and early survival in mosses and ferns (Hepenstrick et al., 2020). These effects may be driven by at least two mechanisms. First, climbing chalk may intensify soil drought stress by further reducing water availability in already shallow cliff substrates. Second, chalk deposition can alter nutrient composition and increase soil pH, potentially affecting organic and inorganic nutrients present in the soil and plant tissues such as nitrogen, carbon, phosphorus, calcium, and magnesium, among others (Barker & Pilbeam, 2015). Because microbial communities can be sensitive to changes in pH, nutrient availability, and ionic composition (Leff et al., 2015; Glassman et al., 2017; Zhang et al., 2024), even small shifts in soil chemistry may disrupt soil- and root-associated microbial assemblages, yet this question needs to be addressed in natural cliff systems.

Cliff plant communities are typically composed of both habitat specialists and generalist species, generating functional mosaics that influence ecological interactions and nutrient dynamics through heterogeneous nutrient return via litter decomposition (Smrithy et al. 2025). Climbing may differentially affect these groups, with cliff specialists appearing more tolerant to some climbing-related disturbances (March-Salas et al., 2018). Cliff specialists have evolved unique functional strategies, such as compact growth forms and extreme physiological stress tolerance, that may confer stronger resilience to stressors associated with climbing (Larson et al., 2000; Villadangos & Munné-Bosch, 2024). However, it remains unclear whether their successful persistence also depends on strong, potentially co-evolved microbial associations.

Although extreme environments have traditionally been considered inhospitable to fungal survival (Gostinčar et al., 2023), mutualistic interactions with fungi, such as arbuscular mycorrhizae (AMF), endophytes, and other rhizosphere-associated symbionts, enhance water and nutrient acquisition, mitigate abiotic stress, and modulate plant secondary metabolism (Rodriguez et al., 2009; van der Heijden et al., 2015; Baron & Rigobelo, 2022). In cliffs, chemical interactions between specialist plants and their rhizosphere fungal microbiome may shape fungal recruitment, influence nutrient mobilization, and even facilitate mineral weathering processes, thereby improving nutrient accessibility in nutrient-poor soil substrates (Uroz et al., 2009; Zhao et al., 2025). Indeed, plants from rocky outcrops appear to exert stronger control over fungal communities than over bacteria or other microbial assemblages, resulting in more specific plant-fungal associations (Nielsen et al., 2010; Tkack et al., 2020; Dumas et al., 2024). Also, specific links with rock-inhabiting fungi, symbiotrophs like AMFs, saprotrophs involved in organic matter turnover, or root and foliar endophytes may be more prevalent in cliff-specialist plants (Krah & March-Salas, 2021; Dumas et al., 2024). In contrast, generalist species, often comprising up to 70% of cliff plant communities (*e.g.*, March-Salas et al., 2018), may rely less on highly specific microbial partnerships and instead associate with broader, more flexible microbial assemblages. Differences in related fungal functional guilds between specialist and generalist cliff-dwelling plants may partly explain why these guilds appear to respond differently to climbing pressures. Determining whether fungal community responses to climbing differ between these cliff-related guilds is therefore critical for predicting vegetation community- and ecosystem-level consequences, and, ultimately, guiding management actions.

In this study, we evaluate the direct and indirect effects of recreational climbing on soil-and plant-associated fungal communities in cliff ecosystems across the Iberian Peninsula. To do so, we sampled soil substrate (referred as soil) and roots of both, cliff-dwelling specialist and generalist plants developed in climbed and unclimbed cliffs and conducted detailed analyses on soil and plant-available nutrients. We then assessed fungal community composition, richness, relative read abundances and functional structure in both soils and roots. Specifically, we tested three main hypotheses: (H1) Climbing, through chalk deposition, increases soil pH and alters nutrient composition, particularly magnesium availability and related ionic balances. (H2) Fungal responses to climbing disturbance differ between specialist and generalist cliff plants, potentially altering functional fungal assemblages. (H3) Soil physicochemical changes mediated by climbing restructure microbial communities, affecting fungal richness, composition, relative read abundance, and functional guilds distribution. By integrating soil chemistry and microbial community analyses in cliffs, our study aims to provide the first comprehensive assessment of how recreational climbing may disrupt soil- and plant-associated microbial communities in cliff ecosystems, with implications for the conservation of these unique yet vulnerable systems.

## Materials and Methods

### Field survey

During June 2021, cliffs where sport climbing is practiced were sampled in northern (Calcena, Aragón), central (Patones, Madrid), and southern (Los Vados, Granada) Spain. Areas located within approximately 0.5 m on either side of established climbing routes (*i.e.*, the typical ascent line defined by bolts and rope) were designated as ‘Climbed site’, whereas areas located at least 1m horizontally from both sides of the climbing route were designated as ‘Unclimbed site’ (March-Salas et al., 2018). Plant species occurring on the sampled cliffs were identified and classified as either cliff specialists (*i.e.*, species occurring exclusively or predominantly in cliffs) or generalists (*i.e.*, species commonly found in other systems but also present in cliffs) (Castroviejo, 1986–2012). Whole plant and soil samples were collected from ‘Climbed’ and ‘Unclimbed’ sites, with soil samples being collected beneath both cliff-specialist and cliff-dwelling generalist plant species. Additionally, to assess whether soil abiotic factors and fungal communities were associated with plant presence, soil samples were also collected from control microsites lacking vegetation. Specifically, for each of the three locations, samples from three climbing routes were collected. At each route, eight individual plants were collected with their rhizosphere soil, including four cliff-specialist and four generalist species, distributed between climbed and unclimbed areas. In addition, four bulk soil substrate samples were collected from unvegetated microsites (controls), with two from climbed areas and two from unclimbed areas. Overall, this sampling design yielded a total of 144 soil samples and 108 plants individuals (root samples).

Soil samples were collected into 50 ml Falcon tubes, which were filled as completely as possible, or with most of the available soil, to ensure representativeness, given the limited soil availability within cliff crevices. Prior to collecting each sample, the metal spoon and other material used for soil extraction was cleaned and flame-sterilized to prevent cross-contamination. At least two photographs were taken at each plant and soil sampling microsite to facilitate *ex situ* plant identification and enable subsequent habitat characterization (*e.g.*, relative soil depth or accumulation).

### Soil chemical analysis

Soil samples not designated for DNA sequencing (*see* below) were dried at 40°C for one week. Dried samples were sieved to < 2 mm, and 0.3-1 g of each sample was subsequently milled using the Mixer Mill MM400 (Retsch GmbH, Germany) at 30 Hz for 40 seconds. To prevent cross-contamination, all materials used for sieving and milling were cleaned with compressor air and, when necessary, rinsed with water between samples. The chemical composition of the samples was analysed to determine concentrations of essential elements for plant development, including total phosphorous (P), potassium (K), sulphur (S), calcium (Ca), magnesium (Mg), total carbon (C), organic and inorganic C (TIC), nitrogen (N) and C/N, as well as plant-available K, Mg, Ca, P, and S. Soil pH and electrical conductivity (also called as Salinity; µS/cm) were also measured.

Total C and N concentrations were determined by elemental analysis via thermal combustion and thermal conductivity detection of CO_2_ and N_2_ (Flash 2000 HT Plus, Thermo Scientific, Bremen, Germany). Plant-available K, Mg, Ca, P, and S were extracted using the Mehlich 3 procedure (Mehlich, 1984), which employs a solution containing NH_4_NO_3_, NH4F, HNO_3_, EDTA, and CH_3_COOH, and quantified using inductively coupled plasma optical emission spectrometry (ICP-OES; 5100 VDV ICP-OES, Agilent, Waldbronn, Germany). For total element concentrations, samples were digested using a mixture of HNO_3_, HF, and H_2_O_2_ (4:2:1, v/v/v) in a microwave digestion system (Mars 6, CEM, Kamp-Lintfort, Germany).

Excess HF was complexed with H_3_BO_3_ prior to analysis, and total element concentrations were measured by ICP-OES. Complete recovery of elements was verified using certified reference material (BCR2, Columbia River Basalt). Soil pH and electrical conductivity were measured in a 1:2.5 soil-to-water suspension using < 2mm sieved soil.

Principal Component Analysis (PCA) was used to reduce dimensionality and explore variation in two separate soil nutrient datasets: one containing ten variables representing total nutrient content, and the other including five variables representing plant-available nutrient content (listed in Table S1–S3; Fig. S1). The first two principal components (PC1 and PC2; Table S2–S3) were included in the data analyses.

### DNA extraction and library preparation of the soil and root samples

Two replicates of 1 g of soil were separated from each soil sample into Eppendorf tubes and stored at -20 °C at the end of the sampling day. In addition, two replicates of approximately 100 mg of the fresh root tissue, selecting the thinnest root fragments, were kept in Eppendorf tubes for DNA extraction. DNA from root and soil samples was extracted using the DNeasy PowerSoil Pro Kit (Qiagen). Some modifications were done in order to improve the quality and quantity of the fungal DNA in the extractions. Briefly, samples, together with the solution CD1, were vortexed with a bead beater at 4m/s for 2 minutes, then, 16 μl of lyticase were added following the recommendation of Pierre et al. (2021) and samples were incubated in a thermoblock at 35°C and 500 rpm for 20 minutes. After the incubation, a new round of bead beater was performed and the manufacturer protocol was followed except for the last elution that was done in 70 μl.

ITS amplicon libraries were prepared using a two-step PCR protocol (Method 4 in Glenn et al., 2019). In the first PCR, around 400bp of the ITS gene were amplified with the primer pair ITS1F-ITS2 (Buee et al., 2009) which contained variable-length tags (unique for each sample) and Illumina adapter tails. The PCR conditions included an activation phase at 95°C for 15min followed by 27 cycles of 30s denaturalization at 94°C, 90s of annealing at 50°C, and 90s of extension at 72°C, ending with a final extension at 72°C for 10 min.

Subsequently, the product was cleaned using Sera-Mag SpeedBeads magnetic bead solution, and eluted in 20μl of TEx1 buffer. From this elution, 1μl was used as template for the indexing of the second PCR, which added Illumina adapters with barcode combinations unique for each sample. The presence of both variable-length tags and barcodes enables the identification and removal of tag-jumping events occurring during sequencing. This second PCR had the same conditions as the first PCR, except that only 5 cycles were used.

Subsamples of the products were checked on 2.5% agarose gel and pooled equimolarly based on band brightness. Pool was cleaned with Promega Magnetic Affinity Beads and sequenced on Illumina MiSeq v3 lanes (2×300bp) at the Institute of Environmental Sciences of Jagiellonian University (Kraków, Poland).

In each round of extraction and PCR, random samples from each location and sample type (soil and root) were chosen to avoid bias related to the laboratory protocols. In any case, six samples were duplicated to calculate the error rate of the laboratory procedures.

### Bioinformatics

After sequencing, reads were demultiplexed using Ultraplex (Wilkins et al 2021) based on their unique barcode and tags combinations, allowing assignment of each sequence to its corresponding sample and removal of sequences with ambiguous or mismatched indices. From the initial 9,777,642 reads retrieved from the Illumina MiSeq sequencing after demultiplexing, Amplicon Sequence Variants (ASVs) were determined by using the DADA2 v. 1.16 analytical pipeline in R (Callahan et al., 2016). Briefly, ITS primers were removed using cutadapt (DADA2 ITS Pipeline Workflow v 1.8, Martin 2011) and reads were quality-filtered (maximum expected errors = 2, maximum N bases = 0, minimum averaged quality = 2). Sequences were dereplicated, the rate error model was inferred and used to implement the sample inference algorithm without discarding singletons. Forward and reverse reads were merged and chimeric sequences removed. Taxonomic assignment was determined for each ASV against the UNITE database v. 8.2. (Abarenkov et al., 2022), using the RDP algorithm implemented in DADA2 pipeline in R. In order to get Operational Taxonomic Units (OTUs) that resemble species level (Tedersoo et al., 2012), ASVs were clustered into 97% sequence similarity using VSEARCH v. 2.8.1 (Rognes et al., 2016) and the most abundant sequence kept as representative of the corresponding OTU. Afterword, LULU curation tool (Frøslev et al., 2017) was used to remove further sequencing errors, resulting in a total of 11,061 OTUs and 6,349,216 reads.

Fungal functional guilds were determined by matching assigned genera to OTUs and the genus-guild database provided by Põlme et al. (2021). From this analysis, 3,490 OTUs were assigned to a primary guild (*sensu* Põlme et al. 2021), which resulted in 4,473,430 guild-assigned reads (31.9% of OTUs and 70.5% of reads). Based on their primary ecological function, assigned guilds were aggregated into three major functional guilds: saprotrophs, pathogroups and symbiotrophs. Lifestyles not directly related to plant or soil cliff environments or unclassified guilds were excluded. Later, the same criteria were applied to define more specific functional guilds: AMF, ectomycorrhizal fungi (ECM) and foliar endophytes in soil samples, while AMF, ECM, foliar endophytes, plant pathogens, and root-associated endophytes in root samples.

Prior to downstream analyses, standardization was applied to samples. Based on the distribution of sequencing depth across samples, six low-depth samples were excluded. The remaining samples were rarefied to a standardized depth of 340 reads per sample using the *rrarefy* function from the *vegan* v2.7.2 package (Oksanen, 2015) in R. This approach accounts for sampling biases and ensures comparability across samples. Rarefaction resulted in the retention of 15.3-53.3% of observed OTU richness per sample and strong conservation of community structure (r = 0.91).

### Estimation of fungal diversity and relative read abundance

Alpha diversity was quantified using Hill numbers (Hill, 2017), which provide a unified framework for diversity estimation. As the order of diversity (*q*) increases, greater weight is assigned to more abundant taxa. Specifically, *q* = 0 corresponds to species richness while *q* = 1 and *q* = 2 correspond to the exponential of Shannon entropy and the inverse Simpson index, reflecting the effective number of common and dominant taxa, respectively (Chao et al., 2014). Hill numbers were calculated from the rarefied OTU matrix using the *specnumber* and *diversity* functions implemented in the *vegan* package. For beta diversity estimation, pairwise dissimilarity matrices were calculated using the Hill numbers framework (*q* = 0, 1, 2) with the *hill_taxa_parti_pairwise* function from the the *hillR* v0.5.2 package (Li et al., 2014).

Furthermore, although metabarcoding read counts are influenced by amplification and sequencing biases, they can provide semi-quantitative estimates of relative taxon abundance and were therefore interpreted as relative rather than absolute measures (Valentini et al., 2016; Lamb et al., 2019; Deagle et al., 2019). Because libraries were pooled equimolarly prior to sequencing and samples were rarefied to an equal sequencing depth, OTU read counts were considered comparable across samples and used as proxy for estimates of rarefied relative read abundance (for simplicity, hereafter referred to as ‘relative abundance’).

#### Statistical analysis

All statistical analyses were conducted in R v4.2.3 (R Core Team, 2023) and results were visualized using *ggplot2* v4.0.1 (Wickham, 2016).

#### Modelling effects on soil composition

Linear mixed-effect models (LMMs) were fitted using the *lme4* v1.1.38 package (Bates et al 2015) to test whether each soil property differed for differences in soil properties across locations, between climbed and unclimbed sites and among cliff-dwelling plant types (here in advance shown as ‘Plant type’). Plant type (cliff-specialist, generalist, unvegetated, the latter named as control), location (Calcena, Patones, Los Vados), climbing effect (climbed or unclimbed) and their two- and three-way interactions were included as fixed effects in the model while soil variables were the response variables, each in separate models. Soil variables included pH, salinity, and nutrient concentrations, specifically total contents of Mg, Ca, P, S, C, and inorganic C as well as plant-available Mg and Ca in soil. Climbing routes nested within locations were added as random intercepts. Random effects were visually inspected to ensure their contribution to model fit. Model assumptions were assessed using simulated residual diagnostics implemented in the *DHARMA* v0.4.7 package (Hartig & Hartig, 2017), including tests for uniformity, dispersion, zero inflation, and residual patterns. The significance of fixed effects was assessed using an analysis of variance (ANOVA) with the *car* v3.1.3 package (Fox et al, 2007). In all statistical models with significant interactions or fixed effects with three factor levels, a *post hoc* test was calculated using the *emmeans* v2.0.1 package (Lenth & Piaskowski, 2025).

The effects of explanatory variables (plant type, location, climbing effect) and their interactions on soil composition were also evaluated following a multivariate approach. A permutation multivariate analysis of variance (PERMANOVA) was implemented in the *vegan* v2.7.2 package (Oksanen, 2015) was conducted on a Euclidean distance matrix based on a set of soil nutrients (those included in PC1 and PC2 of the PCA analysis; Table S2) and pH and salinity. Plant type, location, and climbing status were included as predictor variables. Models were fitted using the *adonis2* function with 999 permutations and predictors were sequentially evaluated (*by = “terms”*). Significant effects were further explored by pairwise comparisons between levels of location and plant type with the *pairwiseAdonis* v0.4.1 package (Martínez-Arbizu, 2017).

#### Modelling effects on fungal community composition

Soil and root alpha diversity and OTUs relative abundance were modelled using generalized linear mixed-effects models (GLMMs) with Poisson error distributions or LMMs to test differences across locations between climbed and unclimbed sites and among cliff-dwelling plant types. Model assumptions were assessed using *DHARMa*. Plant type, location, climbing effect, and their two- and three-way interactions were included as fixed effects and the climbing route nested within locations was added as random intercept. The significance of fixed effects was assessed using ANOVA.

Hill-based dissimilarity matrices were used for conducting beta diversity analysis with the *vegan* package. PERMANOVA was performed including climbing, plant type, location, and their interaction as main effects. Additionally, a canonical correspondence analysis (CCA) was conducted to evaluate the isolated effects of climbing, cliff-dwelling plant type, location and their interaction on community composition in soil and root microbiomes. The significance of the constrained ordination was assessed using permutation tests with the *cca* function, and predictors were sequentially evaluated (*by = “terms”*).

To assess differences among general functional guilds (saprotrophs, pathotrophs and symbiotrophs) in soil and root samples, their OTU relative abundance and species richness were modelled using a negative binomial model implemented in the *glmmTMB* v1.1.13 package (Magnusson et al., 2017). Climbing effect, plant type, location, and the interaction between plant type and climbing effect were included as fixed effects, while route identity was included as random effect. The significance of fixed effects was assessed via ANOVA type analysis. Model diagnostics were assessed using *DHARMa*. *Post hoc* tests with multiple comparison adjustments (Tukey’s HSD) were performed using *emmeans*. This analytical framework was followed to test effects on specific functional guilds (AMF, ECM, foliar endophytes, plant pathogens and root-associated endophytes).

#### Structural equation modelling (SEM)

To evaluate the net effects of climbing on soil- and root-associated fungal diversity, structural equation modelling (SEM) was performed using the *lavaan* package v0.6-19 (Rosseel, 2012). An initial path diagram was constructed based on hypotheses derived from previous studies (Figure S2; Table S4) and was separately tested for the three orders of Hill numbers.

Additional models were separately evaluated for climbed and unclimbed subsets to investigate the net effects of Mg and Ca concentrations on soil and root fungal alpha diversity (*q* = 0-2).

Prior to model fitting, variables included in the path diagram were assessed for multicollinearity by calculating the variance inflation factor (VIF; threshold of 7) (Table S5) with the *usdm* package v2.1.7 (Naimi et al., 2014). Model selection was guided by multiple goodness-of-fit indices, including the root mean square error of approximation (RMSEA), comparative fit index (CFI), and standardized root mean square residual (SRMR). Models with low RMSEA and SRMR (values close to zero) and high CFI (values close to one) were considered to have the best fit, following the recommendations of Kline (2015).

## Results

### Climbing effect on soil chemical composition across locations and plant types

Overall, climbed and unclimbed routes differed significantly in seven of the ten soil chemical properties across locations and plant types (*see* significant climbing effects in Table 1; Fig. 1). However, the magnitude and direction of these climbing effects strongly depended on both the location and the type of cliff-dwelling plant, as evidenced by significant two- and/or three-way interactions involving climbing for all soil chemical properties except plant-available Mg and TIC (Table 1).

**Table 1.** Statistical results on how soil properties and microbiota composition in soils and roots are mediated by cliff-dwelling plant type (cliff-specialist, generalist, and also unvegetated soil samples as control soils), location (Calcena, Patones or Los Vados), climbing (climbed vs. unclimbed), and their two- and three-way interactions. The table reports: Chi-square (*χ^2^*) values from ANOVA tests for soil chemical parameters, total fungal relative abundance (Soil OTUs; Root OTUs), and fungal diversity indices; *χ^2^* values for ANOVA-CCA on soil and root microbiota composition; and *F* values with partial coefficients of determination (*R^2^*) from PERMANOVA for overall soil chemical parameters and fungal community composition. Significant and marginally significant results are highlighted in bold (*0.01 ≤ *p* < 0.05; **0.001 ≤ *p* < 0.01; \*\*\**p* < 0.001; · 0.05 ≤ *p* < 0.1). Abbreviations: *Plant.* = Plant type, *Loc.* = Location, *Climb.* = Climbing Effect, TIC = Total Inorganic Carbon.

| Response | Predictor |  |  |  |  |  |  |
| --- | --- | --- | --- | --- | --- | --- | --- |
|  | *Plant.* | *Loc.* | *Climb.* | *Plant. × Loc.* | *Plant × Climb.* | *Loc. × Climb.* | *Plant. × Loc. × Climb.* |
| ANOVA results |  |  |  |  |  |  |  |
| pH | 7.84* | 2.38 | 11.58*** | 0.79 | 2.52 | 2.97 | 6.62· |
| Salinity | 0.33· | 97.65*** | 0.15 | 11.83* | 0.18 | 6.49* | 10.60* |
| Total Mg | 6.55* | 274.01*** | 4.76* | 17.68** | 1.77 | 9.78** | 9.74* |
| Total Ca | 3.43 | 129.92*** | 6.45* | 5.50 | 0.57 | 3.63 | 11.05* |
| Plant Mg | 4.30 | 25.42*** | 0.18 | 2.70 | 2.56 | 2.13 | 0.55 |
| Plant Ca | 8.52* | 33.02*** | 2.59 | 10.48* | 7.81* | 4.29 | 2.32 |
| Total P | 6.38* | 11.2** | 5.40** | 4.07 | 6.29* | 4.15 | 6.34 |
| Total S | 3.56 | 0.64 | 10.36** | 2.94 | 2.35 | 6.21* | 3.28 |
| Total C | 3.98 | 869.39*** | 8.29** | 3.09 | 7.47* | 0.50 | 3.95 |
| TIC | 11.88** | 73.66*** | 5.40* | 12.74* | 3.43 | 5.95· | 4.83 |
| Soil OTUs | 0.26 | 1.35 | 0.26 | 2.10 | 1.13 | 0.72 | 0.68 |
| Root OTUs | 5.11* | 8.51* | 2.57 | 3.18 | 0.04 | 2.01 | 0.33 |
| Soil q0 | 5.12· | 46.90*** | 0.25 | 16.36** | 12.28** | 2.64 | 4.07 |
| Soil q1 | 1.59 | 0.26 | 0.01 | 10.55* | 5.34· | 0.73 | 1.52 |
| Soil q2 | 7.93* | 0.48 | 0.03 | 9.75* | 1.27 | 1.25 | 1.57 |
| Root q0 | 0.01 | 3.09 | 0.07 | 5.41· | 0.41 | 6.34* | 0.24 |
| Root q1 | 1.55 | 0.17 | 3.55· | 5.40· | 0.01 | 3.69 | 0.96 |
| Root q2 | 0.51 | 0.04 | 2.52 | 4.28 | 0.01 | 1.58 | 2.11 |
| PERMANOVA results |  |  |  |  |  |  |  |
| Overall soil properties | $R^2 = 0.17$<br>$F = 2.12^*$ | $R^2 = 0.50$<br>$F = 64.23***$ | $R^2 = 0.01$<br>$F = 2.81^*$ | $R^2 = 0.01$<br>$F = 0.73$ | $R^2 = 0.01$<br>$F = 1.21$ | $R^2 = 0.01$<br>$F = 1.03$ | $R^2 = 0.01$<br>$F = 0.57$ |
| Soil fungal composition | $R^2 = 0.01$<br>$F = 1.20$ | $R^2 = 0.20$<br>$F = 16.80***$ | $R^2 = 0.01$<br>$F = 1.29$ | $R^2 = 0.03$<br>$F = 1.23·$ | $R^2 = 0.01$<br>$F = 0.97$ | $R^2 = 0.02$<br>$F = 1.40^*$ | $R^2 = 0.02$<br>$F = 0.92$ |
| Root fungal composition | $R^2 = 0.01$<br>$F = 1.70**$ | $R^2 = 0.14$<br>$F = 8.59***$ | $R^2 = 0.01$<br>$F = 1.49^*$ | $R^2 = 0.03$<br>$F = 1.52**$ | $R^2 = 0.01$<br>$F = 1.08$ | $R^2 = 0.02$<br>$F = 1.38^*$ | $R^2 = 0.02$<br>$F = 1.24·$ |
| ANOVA-CCA results |  |  |  |  |  |  |  |
| Soil fungi | 1.09* | 3.13*** | 1.08* | 1.06* | 1.01 | 1.11** | 1.00 |
| Root fungi | 1.21*** | 2.29*** | 1.07· | 1.16*** | 1.00 | 1.11** | 1.01 |

**Figure 1.**
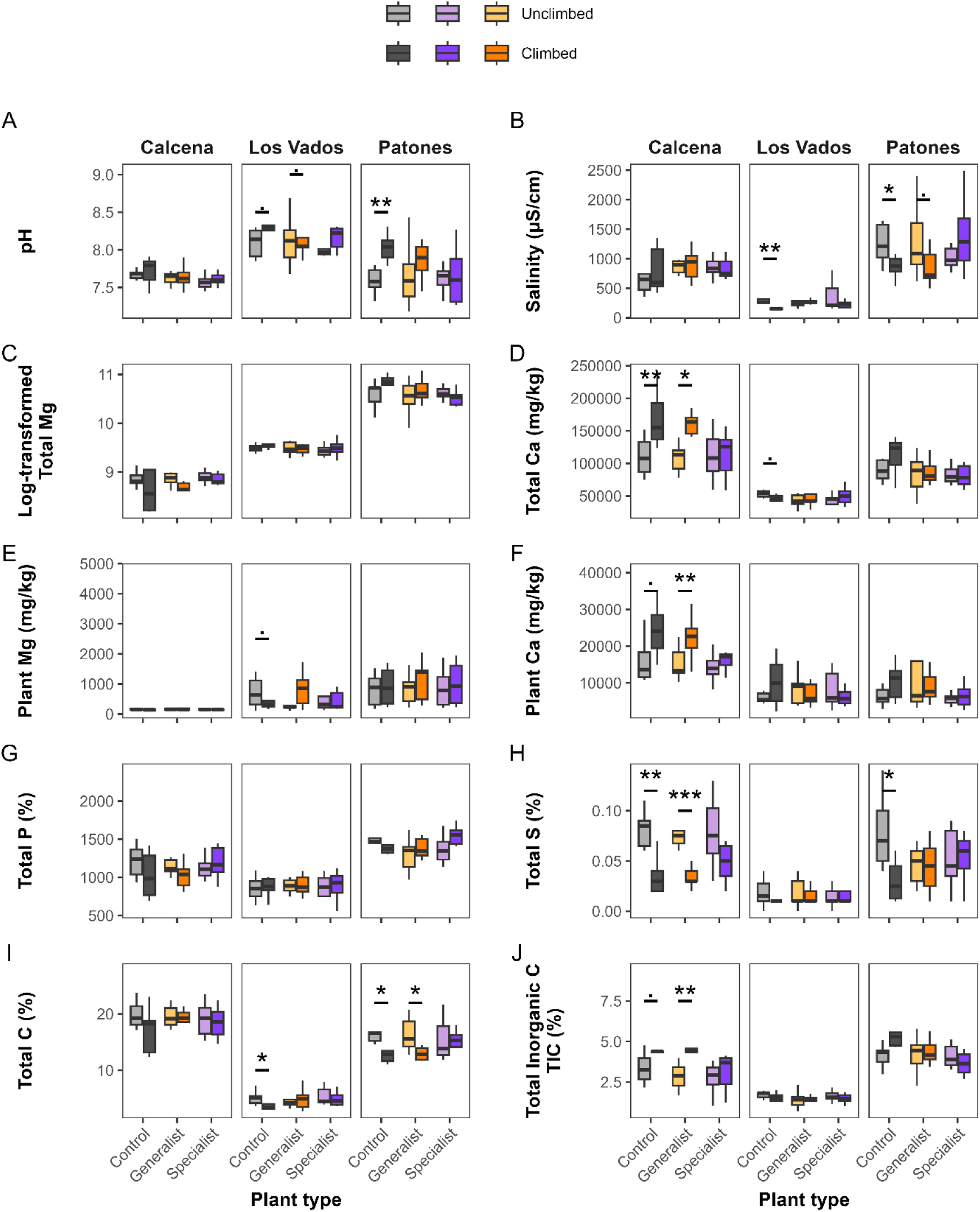
Differences in soil properties between unclimbed (light colours) and climbed (dark colours) sites, differentiated by the cliff-dwelling plant type (cliff-specialist or generalist) present in the sampled soil at each location. Control samples correspond to soils from unvegetated microsites. “*Total*” refers to total element concentrations, whereas “*Plant*” refers to plant-available inorganic fraction of nutrients. Absolute values are showed for all variables except for Total Mg concentrations, originally measured in mg/kg but log-transformed for improved visualization. Asterisks and dots indicate significant and marginally significant differences between climbed and unclimbed sites in each plant type group, respectively (*0.01 ≤ *p* < 0.05; **0.001 ≤ *p* < 0.01; \*\*\**p* < 0.001; · 0.05 ≤ *p* < 0.1).

Soil pH was significantly higher in climbed routes (Table 1), with the strongest differences observed in the control soils from Los Vados and Patones (Fig. 1A). The climbing effect on salinity varied among locations and plant type, as shown in the significant three-way interaction (Table 1). Lower values of salinity in climbed routes were more evident in control and generalist-dominated soils but not in soils with cliff-specialist plants. This trend was found at Patones and Los Vados. The opposite trend, although not significant, was observed at Calcena (Fig. 1B).

The climbing effect on total Mg concentrations varied among locations and plant types, as shown in the significant three-way interaction (Table 1). Total Mg was particularly high in Patones, where the higher concentrations of total Mg in climbed routes were more evident in control and generalist-dominated soils, but not in soils with cliff-specialist plants (Fig. 1C). The climbing effect on total and plant-available Ca varied among locations and plant types, as shown in the significant three-way interaction (Table 1). Total and plant-available Ca showed significant higher concentrations in climbed routes at Calcena, particularly in control and generalist-dominated soils. A similar pattern was observed for TIC (Fig. 1J), while the opposite was true for total S and in total C. Particularly, climbed routes showed lower concentrations of total S in control soils in Calcena and Patones, and in soils with generalist plants in Calcena (Fig. 1H). Total C was significantly lower in climbed routes, particularly in control soils in Los Vados and Patones and in soils with generalistplants in Patones (Fig. 1I). The climbing effect on total P depended on the plant type (Table 1), with control soils located in climbed routes showing lower mean total P concentrations, whereas it was higher in soils with cliff-specialist and generalist plants (Fig. 1J).

PERMANOVA analysis showed that overall soil composition was driven by location, plant type, and climbing as main effects, but their interactions were not significant (Table 1).

### Fungal cliff-dwelling community characterization

After bioinformatics processing, 6,349,216 fungal reads comprising 11,061 OTUs were recovered across the total 252 samples. Following standardization steps, 5,818 OTUs distributed across 244 samples remained for analyses. Specifically, 135 and 109 samples corresponded to soil and root, respectively.

Soil samples contained 3,958 OTUs, of which 615 (15.5%), 210 (5.3%) and 64 (1.6%) were assigned to saprotrophic, pathotrophic, and symbiotrophic functional guilds, respectively. Specific functional guilds included 120 OTUs corresponding to plant pathogens, 21 OTUs to AMFs, 16 to foliar endophytes, 14 to root-associated endophytes and 13 to ECMs. In root samples, 2,550 OTUs were detected. Among these, 175 (6.9%) and 64 (2.4%) were assigned to pathotrophic, and symbiotrophic functional guilds, respectively. Plant pathogens were distributed across 175 OTUs, AMFs across 33, ECMs across 10, and foliar endophytes across 6.

### Climbing effect on fungal diversity and composition across locations and plant types

Climbing activity significantly affected estimates of fungal alpha diversity in soil and root samples, with effects strongly dependent on plant type or location, respectively (Table 1). Specifically, soil fungal species richness (Soil q0) was higher in climbed routes whether soils were colonized by cliff-specialist plants, whereas it was lower in soils with generalist plants and in control soils (Fig. 2). This two-way interaction between climbing and plant type was also significant, although marginally, for soil fungal Shannon index (Soil q1; Table 1).

**Figure 2.**
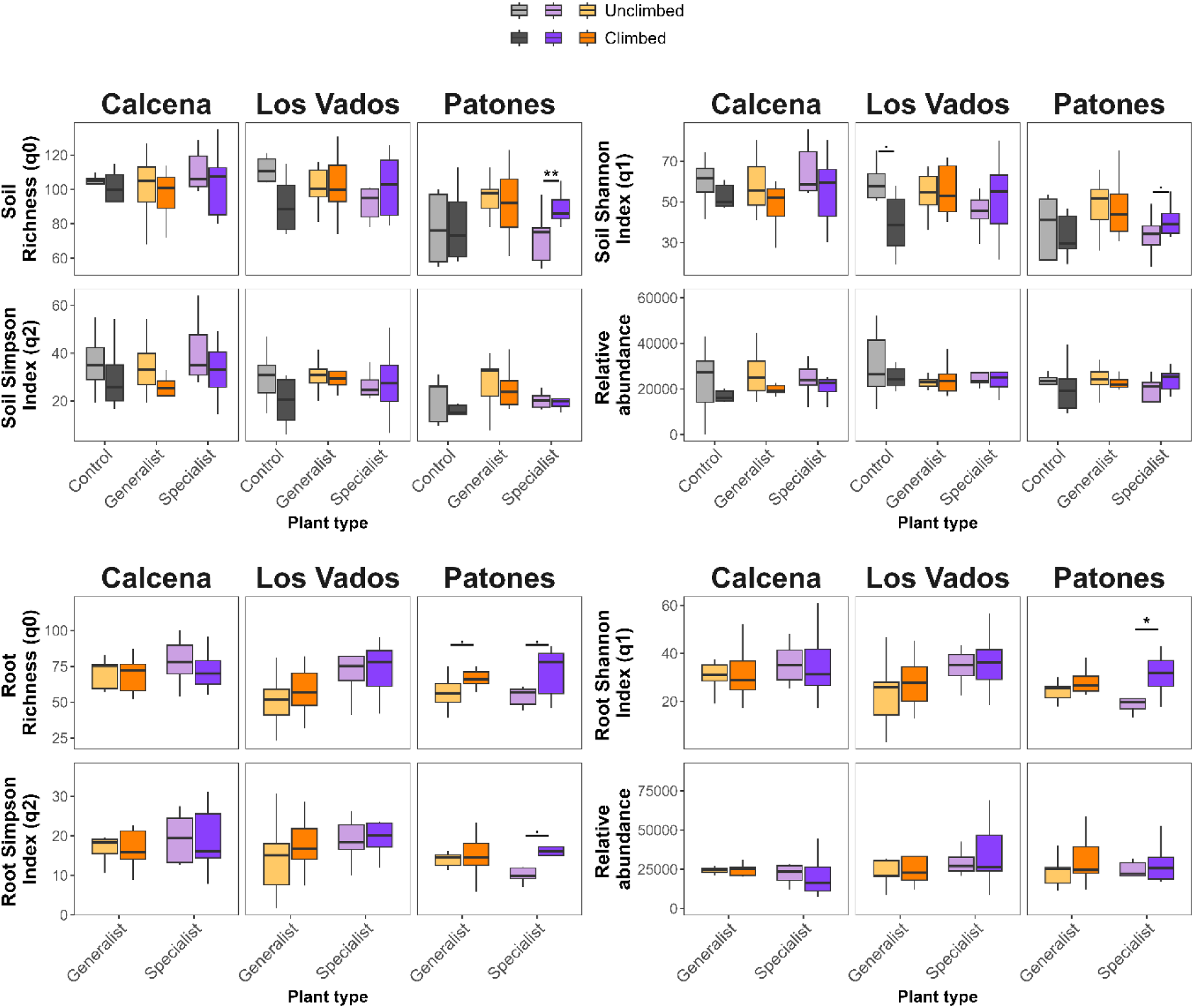
Differences in Hill numbers and OTU relative abundances between unclimbed (light colours) and climbed (dark colours) sites, differentiated by plant type (cliff-specialist or generalist) in (A) soil and (B) root samples. Control samples correspond to soils from sites without vegetation. “*Total*” refers to total element concentrations, whereas “*Plant*” refers to plant-available inorganic fraction of nutrients. Asterisks and dots indicate significant and marginally significant differences between climbed and unclimbed sites in each plant type group, respectively (*0.01 ≤ *p* < 0.05; **0.001 ≤ *p* < 0.01; \*\*\**p* < 0.001; · 0.05 ≤ *p* < 0.1).

Consistent with patterns observed for soil fungal species richness, soils colonized by cliff-specialist plants showed, on average, a higher Shannon diversity in climbed routes, whereas the opposite pattern was observed in soils with generalist plants and in control soils (Fig. 2). The effects between cliff-dwelling plant types depended on the location for both estimates (Table 1), with higher values of both indices in soils with cliff-specialist plants, particularly at Patones.

For root-associated fungal communities, the interaction between climbing and location was significant for fungal species richness (Root q0; Table 1). On average, climbed routes showed higher root fungal species richness at Los Vados and Patones, but lower richness in Calcena (Fig. 2). Climbing also had marginally significant effect on the root fungal Shannon index (Root q1; Table 1), with higher values in climbed routes (Fig. 2). However, significant two-way interactions between the plant type and location for both root fungal species richness and Shannon diversity indicate that higher values of both indexes occurred in the presence of cliff-specialist plants, particularly at Patones (Fig. 2).

Overall, fungal Simpson diversity in soil (Soil q2) and roots (Root q2) was not significantly affected by climbing (Table 1), although root fungal Simpson diversity showed a response pattern similar to that observed for root fungal species richness and Shannon diversity (Fig. 2). In addition to root fungal species richness and Shannon diversity (reported above), the two-way interaction between plant type and location was also significant for soil fungal species richness, Shannon and Simpson diversity. (Table 1; Fig. 2). No factor affected the relative abundance of soil OTUs, whereas the relative abundance of root OTUs was significantly affected by plant type and location, but not by any of their interactions (Table 1; Fig. 2).

Canonical correspondence analysis (CCA) revealed that fungal community composition in both soil and roots was significantly influenced by climbing and by its interaction with location (Table 1). Plant type and its interaction with location also contributed to soil and root-associated fungal community differentiation. Consistent with these results, PERMANOVA showed that climbing led to significantly differentiated soil fungal communities, but this effect depended on location. For root-associated fungal communities, climbing had a main effect but also its effect depended on both plant type and location (Table 1).

### Climbing effect on fungal functional guilds across locations and plant types

At the level of main fungal functional guilds in soil communities (Pathotrophs, Symbiotrophs, Saprotrophs), climbing activity affected the relative abundance of symbiotrophs across locations and cliff-dwelling plant types (Table S6–S7). Particularly, the relative abundance of symbiotrophs was higher in climbed routes in soils colonized by generalist plants (Fig. 3). Climbing effects on OTU richness of pathotrophs differed between the plant types (Table S8), showing higher values in climbed routes when soils were colonized by cliff-specialists (Fig. S3).

**Figure 3.**
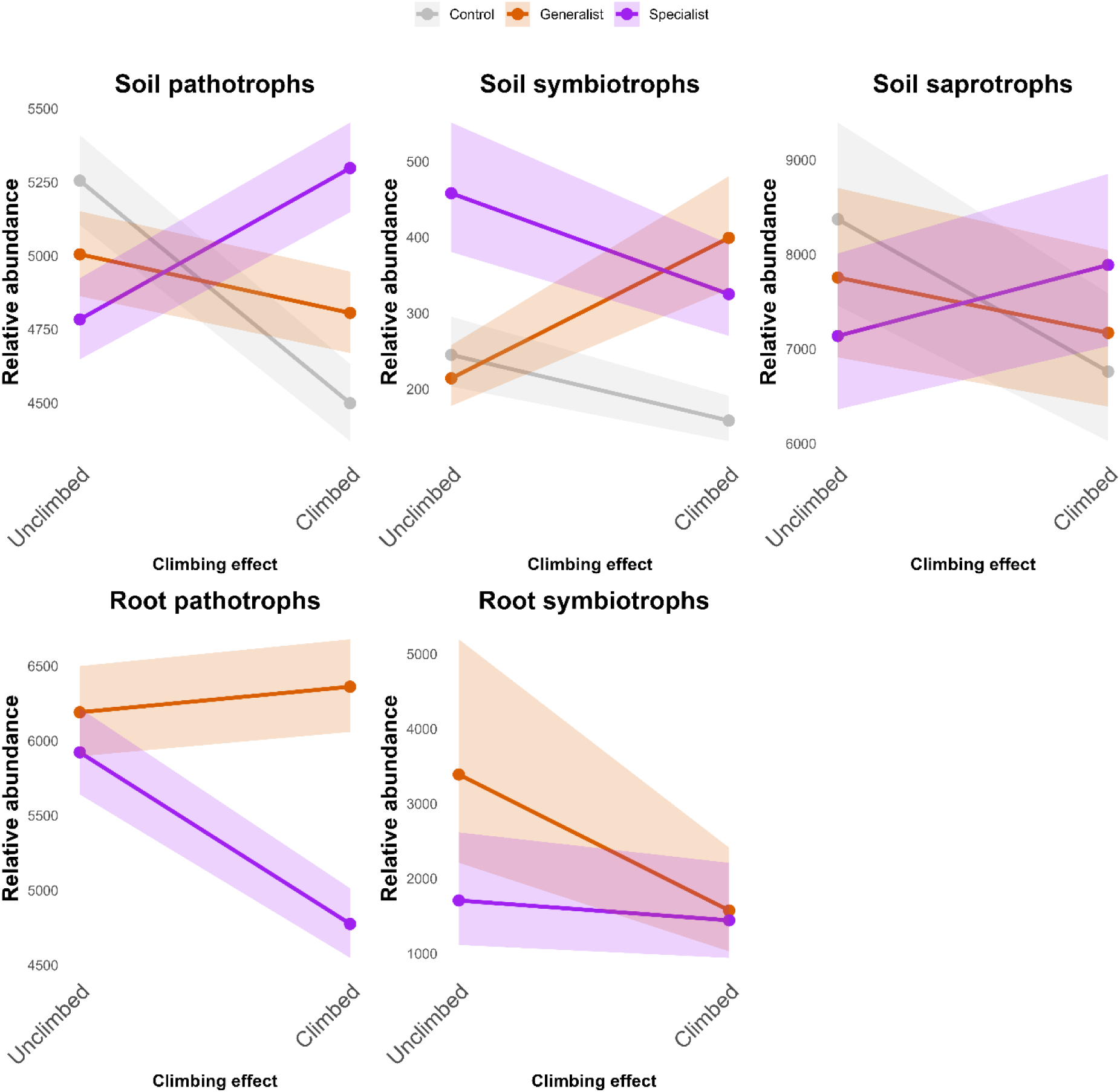
Relative abundance of functional guilds in soil and root samples divided as Pathotrophs, Symbiotrophs and Saprotrophs. Lines represent model-predicted trends with “route” as a random term. Shaded areas indicate 95% confidence intervals. Colours represent the presence of generalist or cliff-specialist plants present in the samples. Grey colour indicates absence of vegetation in the sampled soil. Significance of the slopes are shown in Table S7.

Regarding root-associated main fungal functional (Pathotrophs, Symbiotrophs), climbing activity affected negatively the relative abundance of root symbiotrophs in presence of both plant types (Table S6–S7) but particularly in generalist-dominated soils (Fig. 3). The effect of climbing on the relative abundance of pathotrophs was marginally influenced by both plant type and location as shown by the three-way interaction (Table S6), showing lower values in climbed routes in soils colonized by cliff-specialists (Fig. 3). However, climbing activity positively affected the species richness of pathotrophs in soils in presence of both types of plants but especially in soils dominated by cliff- specialists (Table S8, Fig. S3)

At the level of specific functional guilds in soil communities (AMFs, ECMs, foliar endophytes, plant pathogens), climbing activity affected their relative abundance, with the effect depending on plant type in a couple of functional guilds (Table S7). Specifically, the relative abundance of soil foliar endophytes was higher in climbed routes in soils colonized by cliff-specialist plants, whereas it was lower in soils with generalist plants and in control soils (Fig. S4). The relative abundance of AMFs was increased in climbed routes in soils with both generalist and cliff-specialists and, although not significantly, was decreased in control soils (Table S7, Fig. S4).

Regarding root-associated specific functional guilds (AMFs, ECMs, foliar endophytes, plant pathogens, endophytes), climbing activity decreased the relative abundance of AMFs and ECMs in soils colonized by both cliff-specialist and generalist plants (Table S7, Fig. S6). Climbing activity increased species richness of plant pathogens, especially in soils dominated by cliff-specialists (Table S8; Fig. S7).

### Direct and indirect effects of climbing on soil and root fungal communities inferred through path analysis

A well-fitted structural equation model (SEM; *p* > 0.05; CFI > 0.95; RMSEA < 0.05; SRMR < 0.08) indicated that climbing activity exerted an overall negative effect on both soil- and root-associated fungal alpha diversity through multiple indirect pathways (Fig. 4, S8, S9).

**Figure 4.**
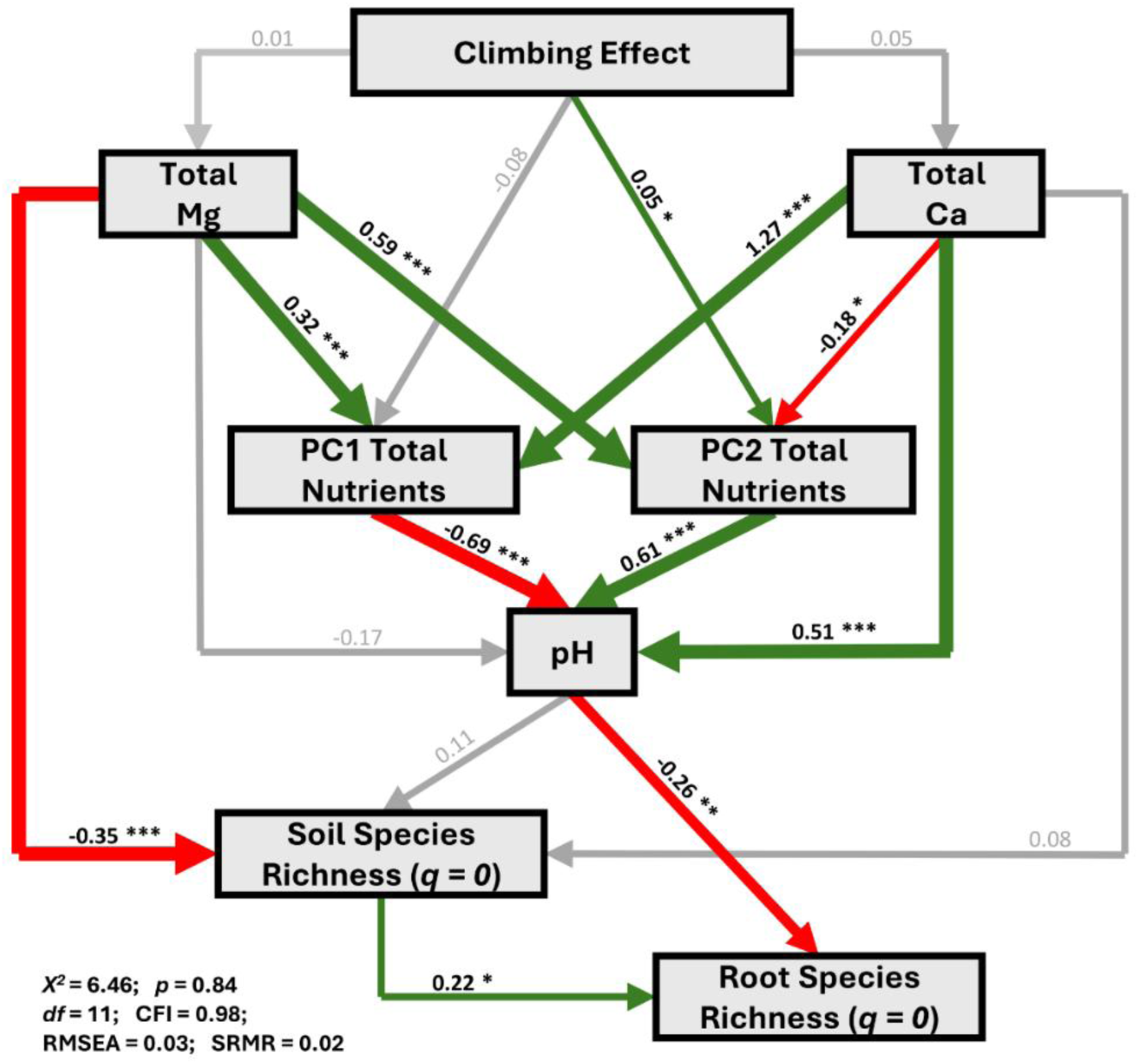
Path diagrams describing the net effects of climbing on soil and root microbiota species richness (*q =* 0), through its direct and indirect influence on soil nutrients and pH. Significant positive and negative relationships are shown in green and red, respectively, while non-significant relationships are shown in grey. Line width represents the strength of significance (*0.01 ≤ *p* < 0.05; **0.001 ≤ *p* < 0.01; \*\*\**p* < 0.001; · 0.05 ≤ *p* < 0.1). Statistics for each structural equation model are shown: Chi-square (*χ^2^*) values; *p* value (*p*); degrees of freedom (*df*); Root Mean Square Error of Approximation (RMSEA); Comparative Fit Index; Standardized Root Mean Square Residual (SRMR).

The most consistent pathway involved an increase in soil pH, which was indirectly driven by climbing via its significant effect on both the second principal component (PC2) of total nutrients and, although not significant, total Ca. Elevated soil pH, in turn, was associated with a reduction in root fungal species richness (Fig. 4). Moreover, total Mg, which increased due to climbing, generated a direct and negative effect on soil fungal species richness, which subsequently translated into lower root fungal richness, considering the positive relationship between soil and root fungal richness. Total Mg also indirectly increased soil pH via PC2 total nutrients (but not by PC1 total nutrients), leading to a significant further decrease in root fungal richness. Overall, these pathway patterns were consistent across the different fungal alpha diversity orders (*q* = 0-2) (Fig. S8).

Several relationships identified in climbed routes were not significant when the SEM model was fitted to the unclimbed dataset. In particular, the positive relationship between soil and root fungal alpha diversity observed in climbed routes was not detected in unclimbed areas (Fig. S9). In contrast, this relationship remained significant in models fitted to the climbed dataset, consistent with a negative effect of climbing and with a significant coupling between soil and root fungal communities in climbed areas. Similarly, the association between soil pH and fungal alpha diversity was significant in the climbed dataset but not in the unclimbed dataset (Fig. S9). Thus, both the soil-root diversity and the pH-fungal diversity relationships were contingent on climbing activity.

## Discussion

Our study shows that recreational climbing alters multiple soil parameters and reshapes the composition of both soil- and root-associated fungal communities across Iberian limestone cliffs. However, the direction and magnitude of these effects varied across locations and strongly depended on the presence and ecological strategy of cliff-dwelling vegetation, since strict cliff specialists seem to hold intrinsic buffering functions against climbing-related impacts on fungi. Our results indicate that climbing activity influences fungal communities through both direct effects and, more importantly, indirect pathways, with shifts in soil pH and other soil properties from the use of climbing chalk acting as a key mediating mechanism. Our insights highlight the sensitivity of plant-associated microbial communities to human disturbances, and how evolved specialist plant-microbial relationships, together with cliff dryness, may help against some of these disturbances. The obtained information can ultimately help guide a sustainable recreational management of cliff ecosystems.

### Climbing as context-dependent driver of cliff soil properties

Overall, our results indicate that climbing activity alters cliff soil properties, although its effects usually vary among sites and the ecological strategies of cliff-dwelling vegetation. Specifically, seven of the ten soil properties showed significant differences associated with climbing, indicating overall chemical effects due to climbing, while nine were significantly influenced by interactions among climbing, location, and/or cliff-dwelling plant type. These predominant context-dependent effects suggest that local abiotic and biotic conditions (*e.g.*, bedrock characteristics, vegetation composition, climate and climbing intensity) modulate cliff soil responses to disturbance. For example, Patones showed frequently pronounced effects, alongside higher average temperatures, lower total precipitation, and increased Mg concentrations and salinity compared to the other locations. Additional factors not directly assessed here may further influence these location-specific responses in soil chemistry. For instance, differences in soil permeability, porosity or chemical transport processes such as drainage or erosion may contribute to location-specific differences by redistributing or diluting nutrient inputs over time (Estrada-Medina et al., 2013). Overall, our results support strong site-specific variability in soil responses, although a generalized influence of chalk deposition from climbers appears evident.

Despite site variability, soil alkalinization emerged as the most consistent effect of climbing activity across all locations, supporting our *first hypothesis*. This pH increase is likely driven by chalk deposition, which supplies carbonate and base cations to cliff soils, while the resulting increase in soil pH acts as a pathway linking climbing disturbance to broader changes in soil chemistry. Additionally, higher Mg and TIC in climbed routes, especially at Calcena, can also be directly attributed to the use of conventional chalk, given its chemical composition. Some commercial chalks also contain calcium carbonate (Ford et al. 2012), which may partly explain increases in Ca in climbed routes. Beyond these direct inputs, elevated Mg concentrations and pH could alter cation exchange dynamics, promoting Ca^2+^ release from soil particles, and enhancing the dissolution and mobilization of Ca-rich particles. Together, these processes not only increase pH and base cations concentrations but also disrupt soil ion balance, thereby affecting overall nutrient mobility and availability.

Consistent with these shifts in soil chemistry, climbed routes showed decreases in salinity and C, S, and P concentrations. Alkalinization may enhance ion mobility and leaching, explaining the reduction in salinity (Butcher et al., 2016). Elevated pH and added Ca could explain reductions in P concentrations through the formation of calcium phosphate (Hopkins & Ellsworth, 2005), while enhanced sulphate mobility under alkaline conditions could account for lower S levels (Moller et al., 2004). The decrease in S and C likely reflects reduced organic matter inputs from cliff plants and soil biota disturbance in climbed routes. Given the critical role of these soil nutrients in plant growth, their reduced availability in climbed routes could be particularly detrimental. In addition, while the effects of climbing on individual soil properties often varied with location and plant type, multivariate analysis revealed a consistent shift in overall soil chemical composition across locations, as indicated by the absence of significant interactions among factors. This pattern suggests that climbing-associated disturbance acts as a directional driver of chemical change in cliff soils.

### Directional and plant-specific effects of climbing on the cliff microbiome

Patterns of fungal diversity and relative abundance across soil and root samples further highlight strong context-dependent effects of climbing disturbance, potentially influenced by plant specialization to cliffs. In soil samples, fungal diversity and relative abundance responded differentially among sites. For example, at Calcena and Los Vados, differences between climbed and unclimbed soils were generally weak or absent, reinforcing the idea that local environmental conditions may modulate the magnitude of climbing impacts on fungal communities, as observed in other cliff-dwelling organisms (Harrison et al., 2022). In contrast, at Patones, when soils were colonized by cliff-specialist plants, species richness and Shannon diversity were higher in climbed areas, suggesting that climbing-related disturbance may, under certain contexts, promote fungal diversity and functional complexity. This is aligned with previous studies reporting increased AMF colonization and total fungal biomass following experimental increments in soil pH (Carrino-Kyker et al., 2016). Further, this pattern is consistent with the ‘intermediate disturbance hypothesis’ (Fox, 1979), suggesting that climbing may enhance habitat heterogeneity and fungal diversity under specific contexts.

Root-associated fungal communities showed higher species richness and Shannon Index in climbed routes, with the strongest climbing differences observed also in Patones. Although an increased environmental stress is often expected to reduce diversity, the observed opposite pattern may be explained by several non-mutually exclusive mechanisms. Climbing activity may increase resource inputs (*e.g.,* chalk-derived particles), selecting for taxa adapted to more alkaline, mineral-rich and dried microhabitats (Elliot et al., 2024), and thereby promoting niche diversification for root-associated fungi. Additionally, disturbance in cliffs may facilitate coexistence of stress-tolerant and opportunistic fungi or root-associated symbiotrophs, while more pathogens and saprotrophs can be found due to decomposing roots. Together, these findings highlight that climbing disturbance affects fungal community structure (Cho et al., 2017), which is also supported by the ‘intermediate disturbance hypothesis’.

Interestingly, the ecological strategies of cliff-dwelling plants appeared to modulate microbial responses to climbing, directly or through their effects on soil composition. While control soils appear to be more susceptible to abiotic and biotic shifts, the presence of vegetation, particularly cliff-specialist species, mitigate these alterations. As shown in soil samples, root-associated fungal diversity of cliff-specialist plants were particularly higher in climbed compared to unclimbed sites, whereas generalist-associated fungal communities showed weaker climbing-related differences. This pattern supports the *second hypothesis*, suggesting that cliff-specialist plants may act as environmental modulators of their associated microbiomes under disturbance, as also observed under other stressors such as drought (Oram et al., 2025). Indeed, cliff vegetation may have acted as an abiotic filter modulating soil matrix composition, likely due to specialized root traits (architecture and exudation), growth strategies or long-term co-adaptation with cliff-associated microbial assemblages (Hartmann et al., 2009; Hodge et al. 2009). Therefore, the strength of this buffering effect mainly depends on plant species-specific traits. Alternatively, cliff-dwelling species persist in nutrient- and water-limited crevices, creating microhabitats that could help isolate fungal communities from disturbance while maintaining soil homeostasis and water retention (Liu et al., 2007; Zhao et al., 2025). Furthermore, cliff-dwelling species enhances stability of fungal communities through slow growth, which reinforces stable root-microbiome associations (Zhao et al., 2025), and low biomass and turnover, which minimize changes in soil composition (De Micco and Aronne, 2012).

### Effects on fungal functional guilds

Patterns in fungal functional guilds reveal that climbing can restructure microbial communities in both soils and roots, although responses differ among plant strategies. In soils, the relative abundance of pathotrophs increased in climbed sites associated with cliff-specialists, whereas declined in soils with generalist plants or without vegetation. This pattern suggests that climbing disturbance may favour fungal pathogens in microsites associated to cliff-specialists, potentially by altering soil chemistry or plant stress. Increased pathogen presence in disturbed environments has been reported in other systems and may reflect shifts in host susceptibility or competitive interactions among microbial guilds (Packer & Clay, 2000; Querejeta et al., 2021; Radujković et al., 2025). A similar pattern was found in the relative abundance of saprotrophs, increasing in climbed sites only in soils with cliff-specialists, highlighting strong plant-mediated modulation of fungal responses. This could be related to the higher magnesium concentrations in climbed routes, which can promote litter decomposition by enhancing microbial metabolic efficiency (Giachetti & Vivanco, 2025). In contrast, soil symbiotrophs, including AMF, tended to increase in climbed sites with generalist plants, while declining or keeping constant in control soils and in soils associated with cliff-specialists. This is consistent with their obligate biotrophic nature and dependence on host plants (Table S9; Meng et al., 2023). Such patterns may indicate increased reliance on mutualistic fungi, particularly AMF, under nutrient imbalance and other climbing-induced stresses (Barea et al., 2011).

Cliffs are frequently mineral-rich and sun-exposed, and retain little water, making them commonly dry. Climbing chalk can further intensifies this dryness, which is known to favour AMF colonization (Cosme, 2023; Oram et al., 2025). The presence and type of cliff vegetation may further alter the associated soil conditions, which could explain the contrasting response. Increased dependence on symbiotic fungi under stressful conditions, such as altered nutrient availability or increased dryness, has been widely documented in plant-microbe systems, as these associations can enhance nutrient acquisition and improve plant tolerance to abiotic stress (van der Heijden et al., 2008; Singh et al., 2011; Acuña-Rodríguez et al., 2020; Zhang et al., 2024). Thus, greater AMF colonization under climbing may reflect higher plant dependency under drought stress, especially for generalist plants not adapted to cliff conditions.

Root-associated communities showed additional contrasts. Root-associated pathotrophs slightly increased in generalist plants under climbed conditions but declined in cliff-specialists, suggesting that plant strategies may modulate microbial responses to disturbance. Root-associated pathotrophs appear to be sensitive to disturbance (Delavaux et al., 2021), but cliff-specialist plants could thereby buffer this effect, supporting to the stabilization of pathotroph communities under climbed routes. Root symbiotrophs, however, declined with climbing across both plant types, with the strongest reduction observed in generalists. This pattern could indicate that disturbance disrupts stable plant-fungal symbioses in roots, potentially through mechanical disturbance, chemical changes in soil, or shifts in plant carbon allocation to microbial partners, as indicated by previous research (Schnoor et al., 2011) and in other disturbed habitats (Radujković et al., 2025). The sensitivity of this disruption may depend on the fungal species involved in the symbiosis, which may explain why these patterns were not always consistent across all locations and plant types (Duan et al., 2011).

The contrasting responses of pathotrophic and symbiotrophic fungi between soil and root compartments may reflect differences in the processes structuring these microbial communities. While soil fungal assemblages are largely shaped by environmental filters such as soil chemistry and disturbance, root-associated communities are more strongly regulated by plant-mediated selection and carbon allocation (Davison et al., 2015). This soil-root decoupling may therefore generate opposite responses of fungal functional guilds under disturbance. Taken together, these results suggest that climbing disturbance does not uniformly affect fungal communities but instead reshapes the balance among functional guilds, although these patterns may vary over time.

### Cascading climbing effects from cliff soil properties to microbiome diversity

Our structural equation modelling (SEM) analyses revealed that climbing activity influences fungal microbiota indirectly through changes in soil properties. This pattern, found in all the measured variables, is in line with our *third hypothesis*, and with studies revealing a tight coupling between microbial communities and soil chemistry (Kristin & Miranda, 2013; Spohn, 2016). Particularly, soil pH has consistently been identified as a major constraint on microbial communities (Lauber et al., 2009; Zhalnina et al., 2015, Naz et al., 2022). The observed increase in soil pH in climbed routes therefore represents a key mechanism underlying the shifts in fungal diversity and composition found in this study.

Soil nutrient content, also altered by climbing, represents another major driver of microbial communities (Kristin & Miranda, 2013, Brabcová et al., 2018; Kang et al., 2021). However, nutrient inputs do not necessarily favour microbial diversity and can even reduce it (Dincă et al., 2022), likely through reductions in susceptible functional guilds or alterations in competitive interactions. Consistently, our SEMs indicate that increased Mg and other alkalinizing nutrients due to climbing negatively affect fungal communities, highlighting soil chemistry as a key intermediary linking climbing to microbes. High Mg concentrations have been associated with decreases in species richness, at least for bacteria (Dash et al., 2025), while evidence for fungi is less common (Yang et al., 2021; Tan et al., 2019 but see Ye et al., 2020). Thus, the negative relationship between Mg and fungal richness shown by our SEMs likely reflects the dominance of Mg-tolerant fungal taxa that outcompete less tolerant species. SEMs also revealed that global effects on soil microbiota are directly translated to root-associated communities, suggesting that root-associated diversity is likely constrained by the surrounding soil microbial pool. This is consistent with the expectation that a large fraction of soil microbiota are formed by root-colonizers (Bulgarelli et al., 2012; Zarraonaindia et al., 2015; Rochefort et al., 2021).

While earlier analyses revealed site-specific and plant-type-dependent responses to climbing disturbance, SEMs highlight a more consistent underlying mechanism linking climbing to changes in fungal communities. In previous models, the effects of climbing varied among locations and plant strategies, reflecting the strong influence of local environmental and biotic conditions. However, SEM results indicate that climbing consistently modifies key soil properties, which negatively affect soil fungal diversity and subsequently propagate to root-associated communities. By integrating additive and cascading effects among soil chemistry and biological responses using SEMs, we reveal how relatively subtle changes in multiple soil parameters can collectively generate stronger and more consistent impacts on fungal communities. Moreover, climbing-induced disturbances may also affect competitive interactions, contributing to the patterns observed (Boddy et al., 2016). Thus, rather than contradicting earlier results, SEMs provide an overall and mechanistic explanation for them.

*To conclude*, our study is the first to assess the impacts of recreational climbing on fungal communities associated in cliff ecosystems. Despite most previous studies focused on the physical disturbance caused by climbing, our results demonstrate that climbing, primarily through chalk use, can also alter the chemical properties of cliff soils and, consequently, reshape the composition of associated fungal communities. Overall, our analyses indicate that these changes can be directional but more likely indirect and/or additive, as climbing chalk application enhances alkalinization, increasing pH and Mg content while modifying soil nutrient composition. The magnitude and direction of these effects can vary across locations and plant ecological specialization, indicating strong fungus-plant and environmental dependencies. This could explain why the impact of climbing is not always consistent, or even overlooked in certain studies (*see* Holzschuh, 2016). However, generalized and cumulative patterns were observed, showing that climbing modifies fungal diversity, community composition, and functional complexity in cliffs. Such changes may result from the loss of taxa sensitive to increased pH or the proliferation of fungal taxa better adapted to altered soil conditions. In some cases, these shifts may also favour microbial taxa that help cliff plants cope with climbing-associated disturbances, including increased soil dryness.

Changes in the presence of cliff-specialist plants may further suggest stronger relationships of these species with specific fungal functional guilds, potentially contributing to explaining their long-term persistence in harsh cliff environments. Understanding the extent of climbing-induced impacts in fungal communities is essential given their key role in nutrient dynamics and plant performance, and thus in cliff ecosystem functioning and resilience.

## Author contribution

MMS, with support from FK, JL, and ALG conceived and designed the study and the field sampling protocol. JL and MMS identified the plant species, together with the field technicians and botany experts. MMS processed the soil and plant samples and organized the chemical analyses. GP and MB performed DNA extraction and library preparation, and GP, FK, and ALG conducted bioinformatic processing of the sequencing data. AGM, with support from MMS, conducted the data analyses and visualization. AGM and MMS wrote the original draft of the manuscript. All authors contributed to discussion, revision, and approved the final version.

## Data availability

Soil chemical data are be deposited in a public repository (10.5281/zenodo.20187505). ITS sequence data are deposited in GenBank under BioProjects PRJNA1466599 (soil samples) and PRJNA1466739 (root samples), and will be public upon publication.

## Acknowledgements

We thank María Urieta and Indradatta deCastro-Arrazola for conducting the fieldwork; Adrián Escudero (IICG-URJC) and Ralph Mangelsdorff (Goethe University Frankfurt) for their help with plant identification; and André Velescu (Karlsruhe Institute of Technology, Karlsruhe, Germany) for conducting the soil and plant chemical analyses.

