## Supplementary Material for "Recreational climbing alters cliff soil chemistry and plant-associated fungal communities"

**Supplementary tables**

**Table S1.** Summary of geographical, environmental and geological characteristics of the study sites. Mean ± Standard Error values and the corresponding units of all soil parameters included in this study are shown. “*Total*” refers to total element concentrations, whereas “*Plant*” refers to plant-available inorganic fraction of nutrients.

| Location | Calcena | Los Vados | Patones |
| --- | --- | --- | --- |
| No. Samples | 81 | 80 | 85 |
| Region | Aragón | Granada | Madrid |
| Altitude (m a.s.l.) | 840 | 170 | 790 |
| Longitude | -1.72 | -3.54 | -3.44 |
| Latitude | 41.65 | 36.79 | 40.89 |
| Rock type | Limestone | Limestone | Limestone |
| Annual mean temperature (°C) | 11.5 | 17.3 | 18.0 |
| Annual precipitation (mm) | 482 | 416 | 276 |
| pH | 7.62 ± 0.02 | 8.11 ± 0.03 | 7.72 ± 0.04 |
| Salinity (µS/cm) | 859.97 ± 40.51 | 262.46 ± 18.77 | 1140.60 ± 62.28 |
| Total N (%) | 1.30 ± 0.03 | 0.27 ± 0.01 | 1.15 ± 0.05 |
| Total C (%) | 18.92 ± 0.50 | 4.75 ± 0.19 | 15.12 ± 0.48 |
| Total inorganic C (%) | 3.42 ± 0.20 | 1.54 ± 0.06 | 4.32 ± 0.19 |
| Total organic C (%) | 15.49 ± 0.59 | 3.21 ± 0.17 | 10.71 ± 0.55 |
| Total C/N | 11.93 ± 0.34 | 11.57 ± 0.23 | 9.31 ± 0.14 |
| Total S (%) | 0.06 ± 0.01 | 0.01 ± 0.01 | 0.05 ± 0.01 |
| Total Ca (mg/kg) | 126341.51 ± 6573.01 | 49684.18 ± 2420.86 | 89102.27 ± 3822.93 |
| Total K (mg/kg) | 9919.54 ± 419.65 | 13220.77 ± 179.57 | 9737.81 ± 344.36 |
| Total Mg (mg/kg) | 6863.27 ± 233.46 | 13181.51 ± 207.83 | 42223.48 ± 1551.37 |
| Total P (mg/kg) | 1128.89 ± 32.10 | 884.91 ± 19.41 | 1416.50 ± 41.31 |
| Plant Ca (mg/kg) | 17936.18 ± 872.36 | 7922.19 ± 687.57 | 8511.92 ± 999.26 |
| Plant K (mg/kg) | 202.93 ± 10.84 | 202.48 ± 40.52 | 242.22 ± 23.38 |
| Plant Mg (mg/kg) | 152.40 ± 5.69 | 628.58 ± 106.63 | 908.37 ± 82.25 |
| Plant P (mg/kg) | 26.59 ± 1.99 | 60.17 ± 16.66 | 78.01 ± 11.13 |
| Plant S (mg/kg) | 44.39 ± 3.15 | 41.10 ± 6.27 | 43.57 ± 3.32 |

**Table S2.** Contributions (*loadings*) of the total soil elements concentrations for the following principal components.

| Variable | PC1  (50.9%) | PC2  (22.3%) | PC3  (15.3%) | PC4  (5%) | PC5  (2.9%) | PC6  (2%) | PC7  (1%) | PC8  (0.4%) | PC9  (0.1%) | PC10  (<0.1%) |
| --- | --- | --- | --- | --- | --- | --- | --- | --- | --- | --- |
| Total N | 0.409 | -0.059 | 0.266 | 0.016 | -0.079 | -0.300 | -0.226 | 0.008 | 0.780 | <0.001 |
| Total C | 0.426 | -0.156 | 0.067 | -0.018 | 0.067 | -0.193 | -0.201 | -0.143 | -0.382 | 0.734 |
| Total Inorganic C | 0.320 | 0.271 | -0.424 | 0.115 | -0.061 | 0.220 | -0.125 | -0.727 | 0.045 | -0.175 |
| Total Organic C | 0.392 | -0.247 | 0.189 | -0.051 | 0.092 | -0.275 | -0.192 | 0.033 | -0.440 | -0.655 |
| Total C*/*N | -0.037 | -0.581 | -0.198 | -0.287 | 0.633 | 0.279 | -0.043 | -0.111 | 0.210 | <0.001 |
| Total S | 0.330 | -0.126 | 0.399 | -0.178 | -0.347 | 0.734 | 0.136 | 0.063 | -0.029 | <0.001 |
| Total Ca | 0.311 | -0.072 | -0.450 | 0.564 | 0.046 | 0.228 | -0.128 | 0.551 | 0.023 | <0.001 |
| Total K | -0.333 | 0.068 | 0.424 | 0.401 | 0.193 | 0.250 | -0.649 | -0.136 | -0.043 | <-0.001 |
| Total Mg | 0.121 | 0.559 | -0.127 | -0.573 | 0.207 | 0.114 | -0.402 | 0.330 | -0.021 | <0.001 |
| Total P | 0.247 | 0.400 | 0.331 | 0.245 | 0.610 | 0.037 | 0.484 | -0.020 | <-0.001 | <0.001 |

**Table S3.** Contributions (*loadings*) of the plant-available soil elements concentrations for the following principal components.

| Variable | PC1  (67.1%) | PC2  (24.6%) | PC3  (4.4%) | PC4  (2.6%) | PC5  (1.2%) |
| --- | --- | --- | --- | --- | --- |
| Plant Ca | -0.062 | 0.973 | 0.210 | 0.057 | 0.009 |
| Plant K | 0.963 | 0.066 | -0.089 | 0.176 | -0.165 |
| Plant Mg | 0.891 | -0.268 | 0.323 | -0.164 | -0.039 |
| Plant P | 0.961 | -0.130 | 0.023 | 0.163 | 0.175 |
| Plant S | 0.839 | 0.431 | -0.251 | -0.210 | 0.031 |

**Table S4.** Literature-based hypothesized mechanisms underlying each path (number) included in the a priori structural equation model illustrated in Figure S1. Full references are included below the table.

| **#** | **Justification** | **Reference** |
| --- | --- | --- |
| **1** | Climbing chalk (MgCO_3_) releases calcium into the soil | Ford, 2012 |
| **2** | Climbing chalk (MgCO_3_) releases magnesium into the soil | Ropp, 2013 |
| **3, 4** | Climbing chalk (MgCO_3_) can disrupt ionic balance in soils | Deru et al., 2023 |
| **5, 7** | Changes in Ca can disrupt ionic balance in soils | Deru et al., 2023 |
| **6, 8** | Changes in Mg can disrupt ionic balance in soils | Deru et al., 2023 |
| **9** | Changes in Ca affect soil microbiota | Li et al., 2025 |
| **10** | Changes in Mg affect soil microbiota | Yang et al., 2021 |
| **11** | Calcium cations increase pH | Ropp, 2013; Hepenstrick et al., 2020 |
| **12, 13** | Cations balance in PC1 and PC2 can affect pH | Haynes and Naidu, 1998 |
| **14** | Magnesium cations increase pH | Ropp, 2013; Hepenstrick et al., 2020 |
| **15, 16** | pH affects microbiota survival | Rousk et al 2010; Zhang et al 2016 |
| **17** | Microbiota composition in soils affect those found associated with roots | Kristin & Miranda 2013 |

**Table S5.** Variance Inflation Factors (VIF) of the variables included in structural equation models (all VIF < 7) after excluding highly correlated variables according to multicollinearity tests.

| Variable | VIF |
| --- | --- |
| Total Ca | 2.43 |
| Total Mg | 4.92 |
| PC1 Total Nutrient | 5.27 |
| PC2 Total Nutrient | 4.49 |
| pH | 2.89 |
| Soil Species Richness (*q = 0*) | 1.36 |
| Root Species Richness (*q = 0*) | 1.18 |
| Soil Shannon Index (*q = 1*) | 1.18 |
| Root Shannon Index (*q = 1*) | 1.14 |
| Soil Simpson Index (*q = 2*) | 1.12 |
| Root Simpson Index (*q = 2*) | 1.12 |

**Table S6.** Statistical results for how the abundance of main functional guilds in soils and roots are affected by cliff plant type (cliff-specialist, generalist, or absence of vegetation), location (Calcena, Patones or Los Vados), climbing effect (climbed vs. unclimbed), and their two- and three-way interactions. The table reports Chi-square (*χ^2^*) values and significant and marginally significant results are highlighted in bold (*0.01 ≤ *p* < 0.05; **0.001 ≤ *p* < 0.01; ****p* < 0.001; · 0.05 ≤ *p* < 0.1). Abbreviations: *Plant.* = Plant type, *Loc.* = Location, *Climb.* = Climbing Effect.

| Response | Predictor | | | | | | |
| --- | --- | --- | --- | --- | --- | --- | --- |
|  | ***Plant.*** | ***Loc.*** | ***Climb.*** | ***Plant. × Loc.*** | ***Plant × Climb.*** | ***Loc. × Climb.*** | ***Plant. × Loc. × Climb.*** |
| Soil Pathotrophs | 0.49 | **18.89***** | 0.01 | 4.36 | 1.20 | 0.71 | 3.70 |
| Soil Symbiotrophs | **6.22*** | 0.02 | **3.78·** | **25.78***** | 0.35 | **8.20*** | 3.28 |
| Soil Saprotrophs | 0.01 | **4.95·** | 0.38 | 1.68 | 1.76 | 2.08 | 1.01 |
| Root Pathotrophs | 2.13 | 0.70 | 0.38 | **5.52·** | 1.20 | 0.55 | **5.02·** |
| Root Symbiotrophs | **11.49***** | **6.72*** | **10.62**** | 2.39 | 0.33 | **6.75*** | 1.46 |

**Table S7.** Statistical results for the how the abundance of main and specific functional guilds in soils and roots are affected by cliff plant type (cliff-specialist, generalist, or absence of vegetation), location (Calcena, Patones or Los Vados), climbing (climbed vs. unclimbed), and the interaction between plant type and climbing effect. The table reports: Chi-square (*χ^2^*) from ANOVA tests for the models and *z*-ratio values from *post-hoc* tests. Significant and marginally significant results are highlighted in bold (*0.01 ≤ *p* < 0.05; **0.001 ≤ *p* < 0.01; ****p* < 0.001; · 0.05 ≤ *p* < 0.1). Abbreviations: *Plant.* = Plant type, *Loc.* = Location, *Climb.* = Climbing Effect.

| Variable | Predictor | | | | *Post-hoc* test | |
| --- | --- | --- | --- | --- | --- | --- |
|  | ***Plant.*** | ***Loc.*** | ***Climb.*** | ***Plant x Climb.*** |  |  |
| Soil Pathotrophs | 0.39 | **17.62***** | 0.01 | 1.31 |  |  |
| Soil Symbiotrophs | **8.20*** | 3.92 | 0.67 | **6.92*** | Control | -0.82 |
|  |  |  |  |  | Generalist | **2.19*** |
|  |  |  |  |  | Specialist | -1.05 |
| Soil Saprotrophs | 0.01 | **4.83·** | 0.39 | 1.85 |  |  |
| Soil AMFs | **6.22*** | 1.07 | **13.98***** | **7.93*** | Control | -0.27 |
|  |  |  |  |  | Generalist | **3.91***** |
|  |  |  |  |  | Specialist | **2.39*** |
| Soil ECMs | **12.80**** | 3.86 | 0.61 | 0.81 |  |  |
| Soil Foliar Endophytes | 3.67 | **9.04*** | **4.27*** | **10.96**** | Control | **-2.55*** |
|  |  |  |  |  | Generalist | 1.33 |
|  |  |  |  |  | Specialist | **-2.66**** |
| Soil Plant Pathogens | 1.05 | **9.05*** | 0.11 | 1.68 |  |  |
| Soil Root-Associated Endophytes | 0.23 | 0.10 | 0.10 | 0.78 |  |  |
| Root Pathotrophs | 2.22 | 0.57 | 0.80 | 0.87 |  |  |
| Root Symbiotrophs | **7.12**** | **17.66***** | **6.79**** | 0.04 | Generalist | **-1.77·** |
|  |  |  |  |  | Specialist | **-1.76·** |
| Root AMFs | **5.44*** | **10.13**** | **4.48*** | 0.42 | Generalist | -0.91 |
|  |  |  |  |  | Specialist | **-1.97*** |
| Root ECMs | 2.46 | **12.16**** | **7.87**** | 0.78 | Generalist | **-2.25*** |
|  |  |  |  |  | Specialist | **-1.9·** |
| Root Foliar Endophytes | 1.68 | 1.86 | 1.04 | 0.98 |  |  |
| Root Plant Pathogens | 0.71 | 0.47 | 0.01 | 0.17 |  |  |
| Root Endophytes | 0.56 | 3.05 | 0.06 | 0.31 |  |  |

**Table S8.** Statistical results for the how the richness of main and specific functional guilds in soils and roots are affected by cliff plant type (cliff-specialist, generalist, or absence of vegetation), location (Calcena, Patones or Los Vados), climbing (climbed vs. unclimbed), and the interaction between plant type and climbing effect. The table reports: Chi-square (*χ^2^*) from ANOVA tests for the models and *z*-ratio values from *post-hoc* tests. Significant and marginally significant results are highlighted in bold (*0.01 ≤ *p* < 0.05; **0.001 ≤ *p* < 0.01; ****p* < 0.001; · 0.05 ≤ *p* < 0.1). Abbreviations: *Plant.* = Plant type, *Loc.* = Location, *Climb.* = Climbing Effect.

| Variable | Predictor | | | | *Post-hoc* test | |
| --- | --- | --- | --- | --- | --- | --- |
|  | ***Plant.*** | ***Loc.*** | ***Climb.*** | ***Plant x Climb.*** |  |  |
| Soil Pathotrophs Richness | 2.84 | **9.19*** | 0.18 | **1.47*** |  |  |
| Soil Symbiotrophs  Richness | 2.37 | **18.67***** | 0.67 | 0.41 |  |  |
| Soil Saprotrophs Richness | 1.44 | **80.40***** | 0.01 | 0.43 |  |  |
| Soils AMFs Richness | 0.85 | 0.65 | 1.57 | 0.32 |  |  |
| Soil ECMs Richness | 0.46 | 0.45 | 0.03 | 1.09 |  |  |
| Soil Foliar Endophytes Richness | 0.99 | 1.33 | 0.03 | 2.15 |  |  |
| Soil Plant Pathogens Richness | 2.03 | **9.70**** | 0.42 | 1.73 |  |  |
| Root Pathotrophs Richness | 1.08 | **5.88·** | **6.50*** | 1.70 | Generalist | 0.77 |
|  |  |  |  |  | Specialist | **2.75*** |
| Root Symbiotrophs Richness | 1.92 | 3.18 | 0.53 | 0.02 |  |  |
| Root AMFs Richness | 0.17 | 1.28 | 0.09 | 1.09 |  |  |
| Root ECMs Richness | 0.21 | 0.88 | 0.83 | 2.62 | Generalist | -0.51 |
|  |  |  |  |  | Specialist | **1.72·** |
| Root Foliar Endophytes Richness | 0.09 | 0.29 | 0.10 | 0.11 |  |  |
| Root Plant Pathogens Richness | 1.07 | **5.88·** | **6.50*** | 1.70 | Generalist | 0.77 |
|  |  |  |  |  | Specialist | **2.75*** |
| Root Endophytes Richness | 0.01 | 0.12 | 0.24 | 0.10 |  |  |

**Table S9.** Comparison between mycorrhizal status, based on Meng et al. (2023), of plant taxa samples in this study and the detection of arbuscular mycorrhiza (AM) by sequencing. OM = obligate mycorrhiza, FM = facultative mycorrhiza, NM = non-mycorrhiza; NA = species not specifically reported.

| **Family** | **Species** | **Mycorrhizal status** | **AM presence** |
| --- | --- | --- | --- |
| Anacardiaceae | *Pistacia terebinthus* L. | OM | Yes |
| Apiaceae | *Petroselinum crispum* (Mill.) Fuss | FM | Yes |
| Asteraceae** | *Jasonia glutinosa* L. | NA | Yes |
| Asteraceae** | *Lactuca serriola* L. | OM | Yes |
| Asteraceae** | *Leontodon* sp. | OM | Yes |
| Asteraceae** | *Pallenis maritima* L. | NA | Yes/No |
| Asteraceae** | *Sonchus tenerrimus* L. | OM | Yes |
| Brassicaceae*** | *Arabidopsis lyrata* (L.) O'Kane & Al-Shehbaz | FM | Yes |
| Brassicaceae*** | *Crambe filiformis* Jacq. | OM | Yes |
| Brassicaceae*** | *Diplotaxis* sp. | OM, FM, NM * | Yes/No |
| Fabaceae | *Anthyllis vulneraria* subsp. *iberica* (W. Becker) Jalas ex Cullen | FM | Yes |
| Fabaceae | *Coronilla valentina* subsp. *glauca* (L.) Batt. | NA (genus-level OM) | Yes/No |
| Fabaceae | *Hippocrepis* sp. | OM | No |
| Fabaceae | *Ononis minutissima* L. | NA (genus-level OM, FM, NM) | Yes/No |
| Geraniaceae | *Geranium lucidum* L. | FM | Yes |
| Lamiaceae | *Salvia Rosmarinus* Spenn. | OM | Yes |
| Lamiaceae | *Teucrium gnaphalodes* L’Hér. | NA (genus-level OM, FM, NM) | Yes/No |
| Lamiaceae | *Teucrium chamedrys* L. | OM | Yes/No |
| Papaveraceae | *Sarcocapnos enneaphyla* L. | NA | Yes/No |
| Plantaginaceae | *Chaenorhinum sp.* | NA (genus-level NM) | Yes |
| Plantaginaceae | *Globularia vulgaris* L. | NA | Yes/No |
| Poaceae** | *Hyparrhenia* sp. | NA | Yes/No |
| Poaceae** | *Melica minuta* L. | OM | Yes |
| Urticaceae | *Parietaria judaica* L. | FM | Yes/No |

** Variable depending on the species
** This family is mostly OM
*** This family is mostly NM, with exceptions.*

**SUPPLEMENTARY FIGURES**

**Figure S1.** Principal Component Analysis (PCA) of (A) total nutrient concentrations and (B) plant-available nutrient concentrations in soil. Direction and length of arrows indicate how strongly each variable contributes to the first two principal components (PC1 and PC2). Colour of cos^2^ values indicates the quality of representation of each variable on the principal component. Summary results from the PCAs are explained below the figure.

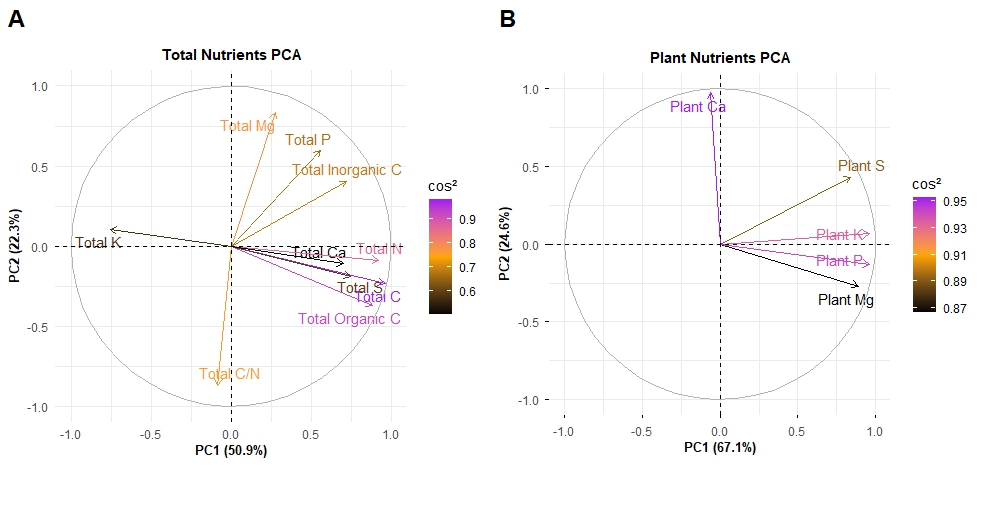

*Variation in total and plant-available soil nutrient concentrations was summarized into two separate principal component analyses (PCAs; Fig. S1). The PCA of total nutrients showed that the first two principal components (PCs) explained a substantial proportion of the total variation in the dataset (PC1 = 50.9%, PC2 = 22.3%). PC1 was positively associated with concentrations of total C and N as well as Ca, S, inorganic and organic C, while concentration of total K showed a negative loading on PC1. PC2 was positively associated with concentrations of total Mg, P and inorganic C while it was negatively associated with the C/N ratio (Fig. S1; Table S2). For plant-available nutrients, the PC1 and PC2 stand for 67.1% and 24.6% of variance, respectively. PC1 was positively correlated with concentrations of plant-available K, P and S. PC2 was positively correlated with plant-available Ca and negatively correlated with plant-available Mg (Table S3).*

**Figure S2.** *A priori* structural equation model used to evaluate the effect of climbing on soil nutrients and on soil and root microbiota diversity. Path numbers correspond to those listed in Table S4.

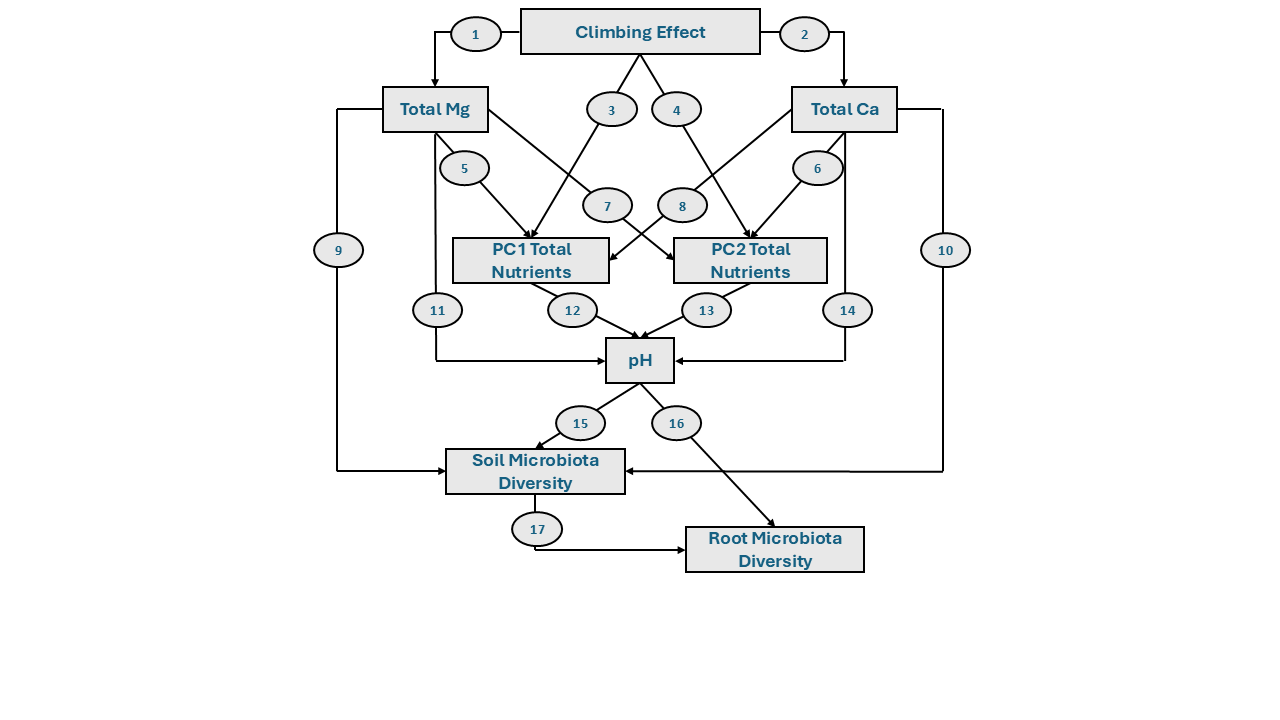

**Figure S3.** Richness of main functional guilds in soil and root. Lines represent model-predicted trends with “route” as a random term. Shaded areas indicate 95% confidence intervals.

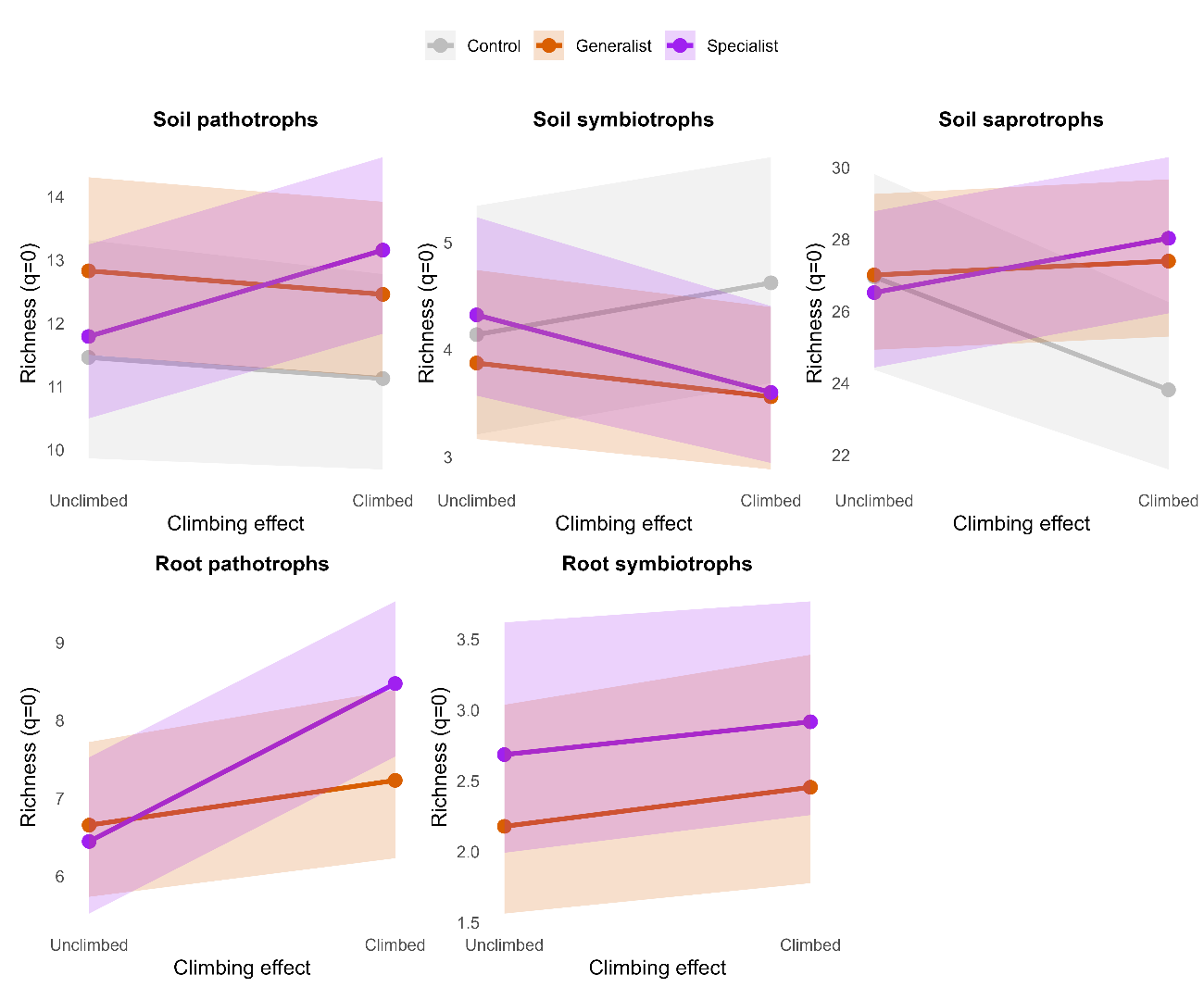

**Figure S4.** Total abundance of specific functional guilds in soil samples divided as arbuscular mycorrhizal fungi (AMF), ectomycorrhizal fungi (ECM), foliar endophytes, and plant pathogens. Lines represent model-predicted trends with “route” as a random term. Shaded areas indicate 95% confidence intervals.

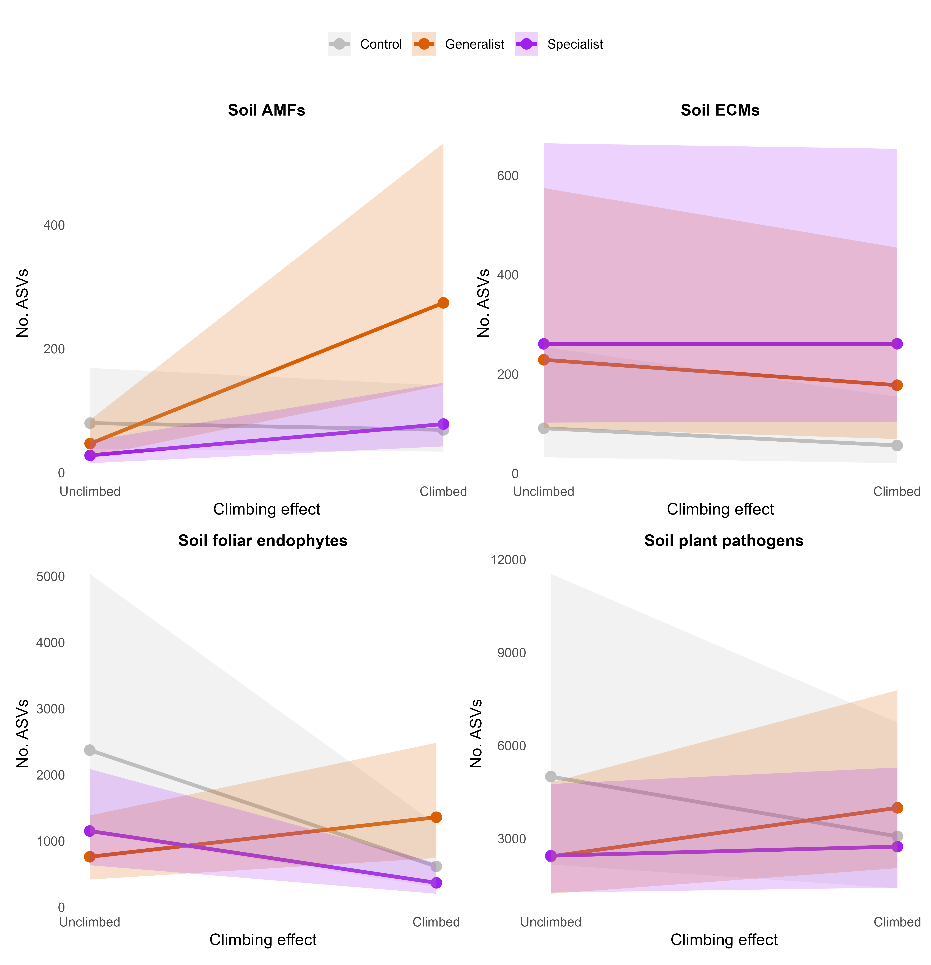

**Figure S5.** Species richness, estimated by Hill numbers (*q* = 0), in specific functional guilds found in soil samples divided as arbuscular mycorrhizal fungi (AMF), ectomycorrhizal fungi (ECM), foliar endophytes and plant pathogens. Lines represent model-predicted trends with “route” as a random term. Shaded areas indicate 95% confidence intervals.

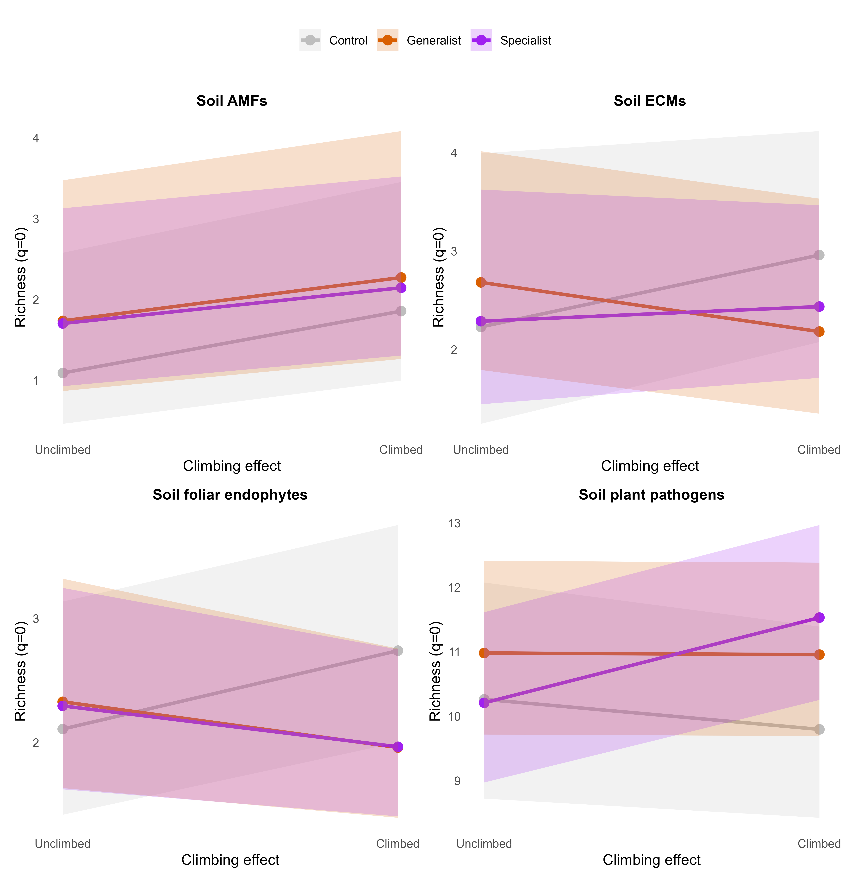

**Figure S6.** Total abundance of specific functional guilds in root samples divided as arbuscular mycorrhizal fungi (AMF), ectomycorrhizal fungi (ECM), foliar endophytes, plant pathogens and root-associated endophytes. Lines represent model-predicted trends with “route” as a random term. Shaded areas indicate 95% confidence intervals.

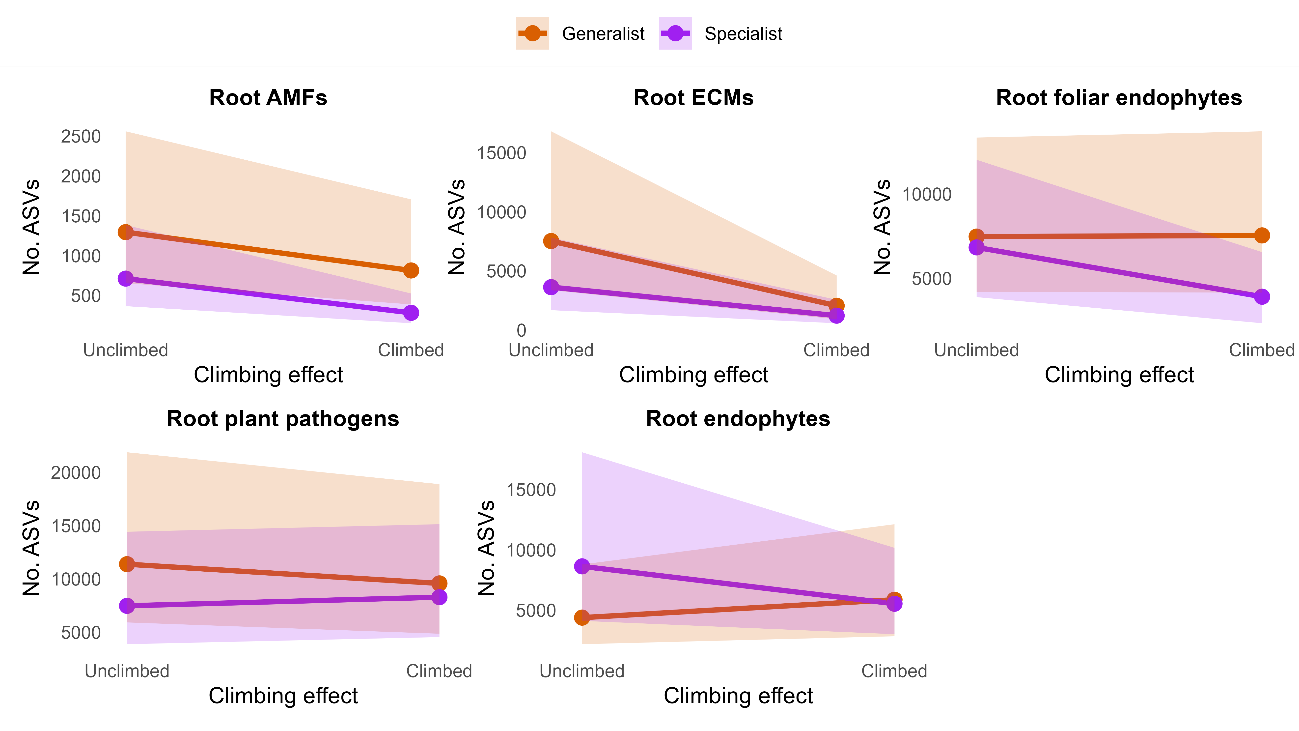

**Figure S7.** Species richness, estimated by Hill numbers (*q* = 0), in specific functional guilds found in root samples divided as arbuscular mycorrhizal fungi (AMF), ectomycorrhizal fungi (ECM), foliar endophytes, plant pathogens and root endophytes. Lines represent model-predicted trends with “route” as a random term. Shaded areas indicate 95% confidence intervals.

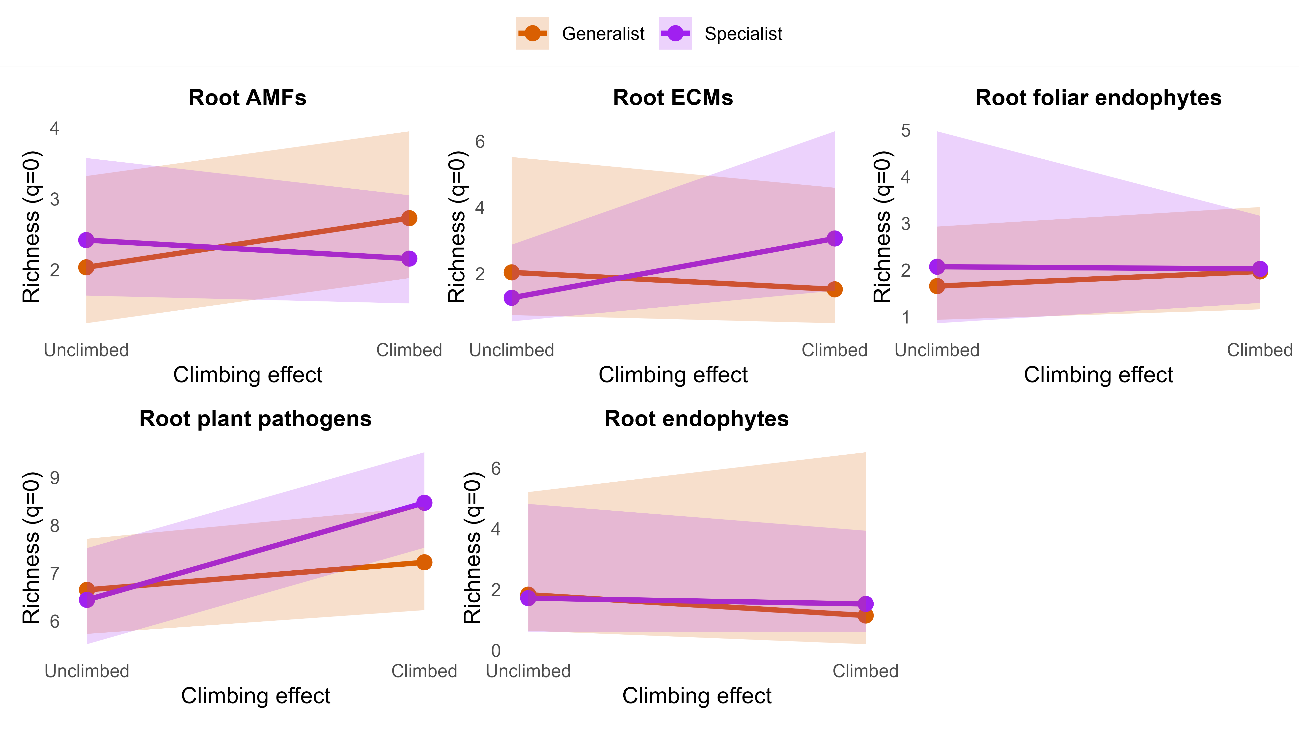

**Figure S8.** Path diagrams describing the net effects of climbing on soil and root microbial diversity, measured using Shannon diversity (*q =* 1) and Simpson diversity (*q* = 2), through its direct and indirect influence on soil nutrients and pH. Significant positive and negative relationships are shown in green and red, respectively, while non-significant relationships are shown in grey. Line width represents the strength of significance (*0.01 ≤ *p* < 0.05; **0.001 ≤ *p* < 0.01; ****p* < 0.001; · 0.05 ≤ *p* < 0.1). Statistics for each structural equation model are shown: Chi-square (*χ^2^*) values; *p* value (*p*); degrees of freedom (*df*); Root Mean Square Error of Approximation (RMSEA); Comparative Fit Index; Standardized Root Mean Square Residual (SRMR).

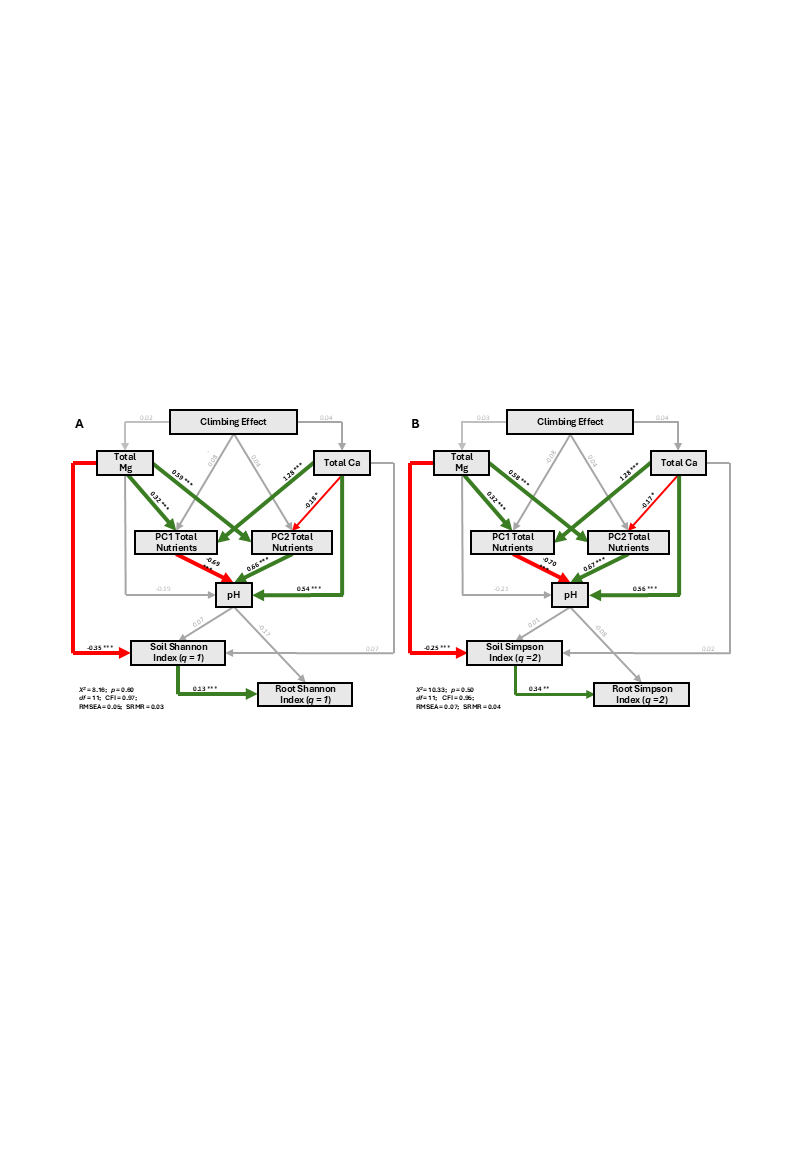

**Figure S9.** Path diagrams describing the net effects of magnesium and calcium concentrations on soil and root microbiota diversity (species richness, Shannon index and Simpson index). Models were fitted separately using data from (A) climbed and (B) unclimbed routes. Significant positive and negative relationships are shown in green and red, respectively, while non-significant relationships are shown in grey. Line width represents the strength of significance (*0.01 ≤ *p* < 0.05; **0.001 ≤ *p* < 0.01; ****p* < 0.001; · 0.05 ≤ *p* < 0.1). Statistics for each structural equation model are shown: Chi-square (*χ^2^*) values; *p* value (*p*); degrees of freedom (*df*); Root Mean Square Error of Approximation (RMSEA); Comparative Fit Index; Standardized Root Mean Square Residual (SRMR).

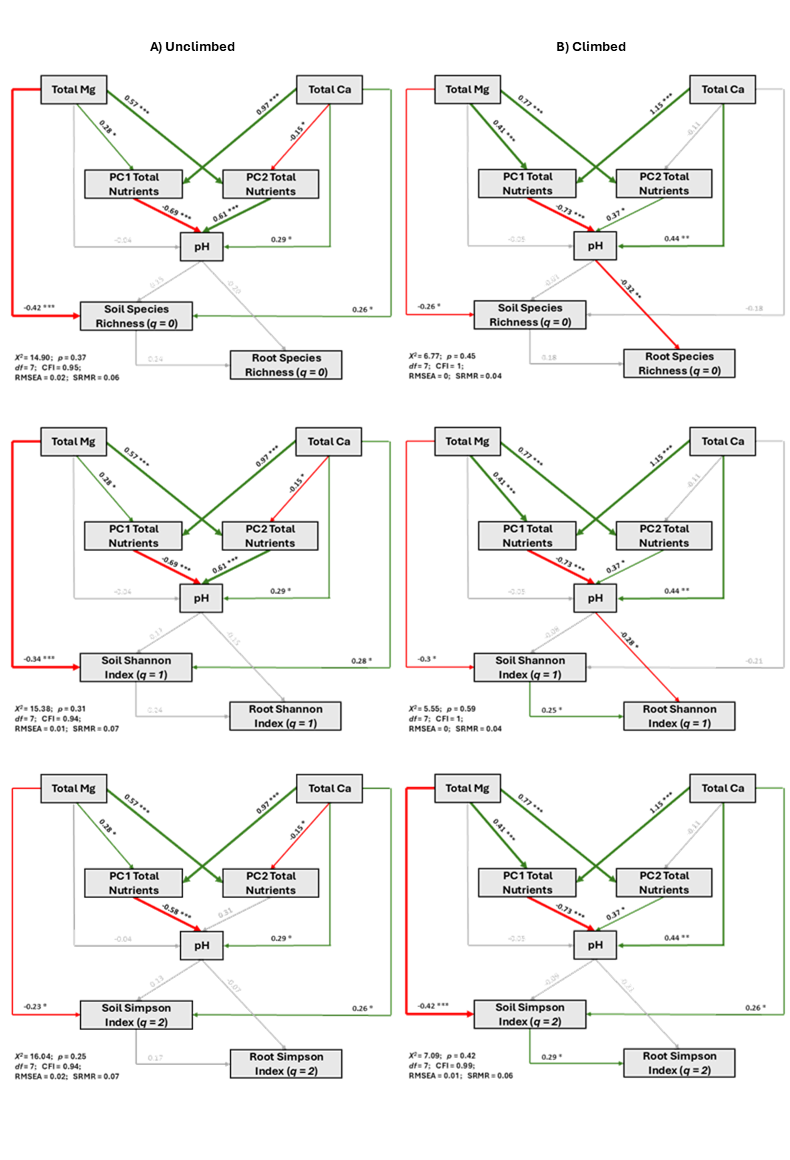
